# CAMKV Kinase Signaling Is a Novel Therapeutic Avenue with Prognostic Relevance in Neuroblastoma

**DOI:** 10.1101/2024.02.19.581040

**Authors:** Yang Yu, Yanling Zhao, Zhongcheng Shi, Feng Cheng, Larry L. Wang, Jong Min Choi, Kan Li, Daniel Silverman, Dan Qi, Jun Wang, Saurabh Agarwal, Brian R Rood, Jeffrey S. Dome, Muller Fabbri, Joanna S. Yi, Erxi Wu, Sung Yun Jung, Chunchao Zhang, Jianhua Yang

## Abstract

Neuroblastoma (NB) can be a highly aggressive malignancy in children. However, the precise mechanisms driving NB tumorigenesis remain elusive. This study revealed the critical role of CREB phosphorylation in NB cell proliferation. By employing a CRISPR-Cas9 knockout screen targeting calcium/calmodulin-dependent protein kinase (CaMK) family members, we identified the CaM kinase-like vesicle-associated (CAMKV) protein as a kinase that mediates direct phosphorylation of CREB to promote NB cell proliferation. *CAMKV* was found to be a transcriptional target of MYCN/MYC in NB cells. CAMKV knockout and knockdown effectively suppressed NB cell proliferation and tumor growth both in vitro and in vivo. Bioinformatic analysis revealed that high CAMKV expression is significantly correlated with poor patient survival. High-risk NB frequently had high CAMKV protein levels by Immunohistochemical staining. Integrated transcriptomic and proteomic analyses of CAMKV knockdown cells unveiled downstream targets involved in CAMKV-regulated phosphorylation and signaling pathways, many of which are linked to neural development and cancer progression. We identified small molecule inhibitors targeting CAMKV and further demonstrated the efficacy of one inhibitor in suppressing NB tumor growth and prolonging the survival of mice bearing xenografted tumors. These findings reveal a critical role for CAMKV kinase signaling in NB growth and identified CAMKV kinase as a potential therapeutic target and prognostic marker for patients with NB.

## Introduction

Neuroblastoma (NB) is the most common extracranial solid tumor in children, accounting for 8-10% of childhood tumors (*1*). Despite its diverse clinical manifestations and variable response to therapy, the prognosis for high-risk NB remains unacceptably low (*1*, *2*). Genomic amplification of the *MYCN* oncogene is associated with tumor aggressiveness and serves as a major prognostic factor in NB (*3–6*). In the absence of *MYCN* amplification, overexpression of MYC characterizes the aggressive phenotype of high-risk NB (*7–9*). However, direct targeting of a transcription factor such as MYCN/MYC is challenging, though some recent reports have shown some hope (*10*). Hence, unraveling other oncogenic signaling pathways is imperative for identifying novel therapeutic targets in NB.

The transcription factor, cyclic AMP response element-binding protein (CREB), is a well-characterized member of the basic leucine zipper super-family. It serving as a critical nidus connecting upstream kinase signaling to downstream target gene expression. CREB activation is mediated through the phosphorylation of Serine 133 in its transactivation domain (*11*, *12*), which facilitates the recruitment of p300/CREB-binding protein (CBP) and subsequent histone acetylation of, along with chromatin remodeling (*13*). CREB promotes cell proliferation by upregulating multiple cell cycle-related genes, including *CCNA1* (*14*), *CCNA2* (*15*), and *CCND1* (*16*). CREB is also necessary for the survival of neuronal cells and NB cells (*17–22*). CREB phosphorylation is known to enhance melanoma’ resistance to chemotherapy and radiotherapy (*23*). However, the exact function and regulatory role of CREB in NB remains unclear.

Calcium/calmodulin-dependent protein kinases (CaMKs) constitute a family of serine/threonine kinases activated by stimuli elevating intracellular calcium levels (*24*, *25*). The CaMK regulatory domain harbors a CaM binding site, and the binding of CaM initiates autophosphorylation to release autoinhibition and activate kinase activity (*26*). Various CaMK family members have been reported to play pivotal roles in cancer development (*27*). Notably, members of the CAMK2 subfamily (CAMK2A, CAMK2B, CAMK2D, and CAMK2G) and CAMK4 have been shown to serve as CREB-activating kinases, which contribute to oncogenesis in diverse tumor types (*28*, *29*).

Intriguingly, CaM kinase-like vesicle-associated protein (CAMKV), a gene structurally akin to the CaMK family but considered a pseudokinase (*30*), has been linked to dendritic spine maintenance (*31*) and activity-dependent bulk endocytosis in neurons (*32*). CAMKV has also been identified as a prognostic biomarker in human endometrial carcinoma (*33*). A recent study revealed *CAMKV* as a transcriptional target of MYCN/MYC in NB (*34*).

In this study, we identified a novel mechanism by which CAMKV controls CREB phosphorylation and activation in NB cells. We demonstrated CAMKV to be a bona fide CREB kinase that mediates CREB phosphorylation and promotes NB cell proliferation. Through integrated transcriptomic and proteomic analyses of CAMKV knockdown cells, we uncovered downstream genes/proteins involved in CAMKV-regulated phosphorylation and signaling pathways. Additionally, we identified small molecule inhibitors that target CAMKV and examined their efficacy *in vitro* and *in vivo*. Overall, our study highlights the role of CAMKV as a critical kinase regulated by MYCN/MYC to promote NB tumor growth and represents a new, druggable NB target.

## 2. Material and methods

### 2.1 Antibodies, chemicals, drugs, and other materials

Antibodies for phospho-CREB (Ser133) (#9198L), CREB (#9104S), mouse IgG (#7076), and rabbit IgG (#7074) were from Cell Signaling Technology; antibodies for CAMKV (D-18) (#sc-102406), CAMKV (2F3-1A2, sc-517082), c-Myc (#sc-40), N-Myc (#sc-56729), CCNA2 (#sc-271682), His-tag (H-3, (#sc-8036), CAMK4 (H-5, c55501) and GATA2 (#sc-267) were from Santa Cruz Biotechnology. Monoclonal Anti-FLAG M2 and Monoclonal Anti-β-actin were from Sigma-Aldrich. Phospho-GATA2 (Ser192) (#PA5-105538) was from Thermo Fisher Scientific. Phosphatase inhibitor cocktail 2 (#P5726) and cocktail 3 (P0044) were from Sigma-Aldrich. Glutathione Sepharose 4B (#17-0756-01) was from Sigma-Aldrich. All small molecule inhibitors used in our assays were listed in table S1.

### 2.2 Cell culture

Human NB cell lines: SK-N-AS, IMR32, and SH-SY5Y were from ATCC; NGP and LAN-1 were from DSMZ; CHLA-255, CHLA-136, and SK-N-BE2 were from Children’s Oncology Group (COG); NB-19 was gifted from Dr. Andrew Davidoff; LAN-6 was gifted from Dr. Robert Seeger. Human embryonic kidney cell line HEK-293T was from ATCC. NGP, SK-N-AS, IMR32, NB-19, CHLA255, LAN-1, LAN-6, SK-N-BE2, and SH-SY5Y were grown in RPMI1640 containing 20% fetal bovine serum (Invitrogen), 100 units/ml penicillin, and 100 µg/ml streptomycin. HEK293T cells were grown in DMEM containing 10% fetal bovine serum. Patient-derived xenograft (PDX) COG-N-519x and COG-N-564x cells were obtained from the COG Cell Culture and Xenograft Repository and maintained in IMDM containing 20% fetal bovine serum, 4 mM L-glutamine, and 1 × ITS (5 µg/mL insulin, 5 µg/mL transferrin, 5 ng/mL selenous acid) (Biotechne, AR013, Minneapolis, MN, USA).

### 2.3 Plasmids

The full-length open reading frame of the wild-type (WT) human *CAMKV* was subcloned into the mammalian expression vector *pcDNA3.1* with a C-terminal FLAG tag. The expression constructs of *CAMKV* mutants were generated by site-directed PCR mutagenesis and verified by DNA sequencing. The lentiviral packing vector *psPAX2* (Addgene plasmid # 12260) and envelope vector *pMD2.G* (Addgene plasmid # 12259) were gifts from Didier Trono. PB-iCas9 was a gift from Xiaojun Lian (Addgene plasmid # 160048). For prokaryotic expression of CREB proteins, cDNA encoding CREB-WT was subcloned into a modified *pGEX* vector to generate the N-terminal GST-tagged fusion proteins. A *Trc2*-lentiviral vector was used to generate shRNA plasmids for *CAMKV*, *CREB,* and *CCNA2*. All target sequences for each gene are described in table S2.

### 2.4 Generation of CRISPR-Cas9 knockout constructs

The protocol for generating a CRISPR-Cas9 library targeting *CaMKs* was described previously (*35*). The lentiCRISPR v2 plasmid was developed by Zhang Lab (Addgene plasmid # 52961) (*36*). Target sequences specific for the 11 CaMK family members were individually cloned into the lentiCRISPR-Cas9 plasmid. The knockout plasmids were transfected into SK-N-AS cells seeded in 6-cm dishes with Fugene 6 and selected with puromycin (0.5 μg/ml) for 5 days. Target sequences specific to the 11 members of CaMKs are listed in table S3.

### 2.5 Generation of inducible shRNA knockdown construct

A doxycycline-inducible TRE-CAMKV shRNA piggyBac transposon vector containing a TRE-sh-CAMKV cassette followed by a phPGK-driven puroR-T2A-rtTA cassette was constructed. This vector was co-transfected with a plasmid expressing the piggyBac transposase into SK-N-AS cells. Cells were selected with neomycin at 500 µg/mL to generate polyclonal cell populations that stably express inducible *CAMKV* shRNA. To induce the expression of CAMKV shRNA, doxycycline was administrated at 100 ng/mL for the indicated time.

### 2.6 Generation of inducible CRISPR-Cas9 knockout constructs

A doxycycline-inducible TRE-Cas9 piggyBac transposon vector containing a cassette of a hPGK promoter-driven puroR-T2A-rtTA cassette was co-transfected with a plasmid containing pCMV-driven piggyBac transposase into CHLA-136-Luc cells. Cells were selected with neomycin at 500 µg/mL. Then a second lentivirus vector (pCDH-BL) containing a doxycycline-inducible TRE-single-guide RNA (sgRNA) of *CAMKV* with a blasticidin (BL) selection marker was transduced into CHLA-136-Luc cells followed by subsequent BL selection to generate polyclonal cell populations that stably express inducible Cas9 and the *CAMKV* sgRNA. To induce the expression of Cas9 protein and sgRNA, doxycycline was administrated at 100 ng/mL for the indicated time.

### 2.7 Immunoblotting

Cell lysates were obtained by lysing cells with RIPA lysis buffer (25 mM HEPES at PH 7.7, 135 mM NaCl, 1% Triton X-100, 25 mM b-glycerophosphate, 0.1 mM sodium orthovanadate, 0.5 mM phenylmethylsulfonyl fluoride, 1 mM dithiothreitol, 10 μg/ml aprotinin, 10 μg/ml leupeptin, 1 mM Benzamidine) containing phosphatase inhibitor cocktail 2 and 3 (Sigma, P5726 and P0044). Supernatants containing proteins were collected after centrifugation at the highest speed for 15 minutes at 4°C, then resolved by SDS polyacrylamide gel electrophoresis and transferred to PVDF membranes. The membranes were then incubated with corresponding primary antibodies in 1 × Tris Buffered Saline with Tween (TBST) containing 5% milk or bovine serum albumin (BSA) overnight at 4°C and horseradish peroxidase-conjugated secondary antibodies for 1 hour at room temperature. Then the membranes were exposed to X-film and visualized using the ECL Western detection system (#32106, Thermo Scientific).

### 2.8 Purification of recombinant GST-CREB protein

The *pGEX-CREB* expression plasmids were transformed into E. *coli* BL-21 pLys (Invitrogen) and selected on the Luria–Bertani (LB) agar plates containing 100 μg/ml ampicillin. The colonies were inoculated and cultured in LB medium containing 100 μg/ml ampicillin at 37°C. Once OD600 reached 0.6-08, isopropyl-b-D-1-thiogalactopyranoside (IPTG) was added at a final concentration of 1 mM to induce expression of GST-tagged CREB for 4 h at 30°C. Bacteria were pelleted and lysed with 1 × extraction buffer (50 mM Tris-HCl, pH 8.5, 100 mM NaCl, 1 mM EDTA, 1 mM DTT, 50 mg/mL lysozyme, 10 μg/mL aprotinin, 10 μg/mL leupeptin, and 1 mM PMSF). The bacteria were sonicated at 4°C in 1% Sarcosyl (Sigma), followed by adding Triton X-100 (1%), 5 μg/mL DNase I, and 5 μg/mL RNase. The lysate was centrifuged at 15,000 x g, and the supernatant containing GST-fusion proteins was collected. GST protein was purified using glutathione Sepharose 4B beads (Sigma) by rotating overnight at 4°C. Beads were then washed three times in extraction buffer containing 0.5% Triton X-100 and one extra wash in extraction buffer containing 0.1% Triton X-100. GST-CREB proteins were eluted in elution buffer (30.7% glutathione, 50 mM Tris–HCl, pH 8.0, 20% glycerol, 5M NaCl) and dialyzed in 1 × PBS. The protein concentrations were then assessed with a Bradford Protein Assay (Bio-Rad). The proteins were visualized by 10% SDS-PAGE and Coomassie blue staining of the gel.

### 2.9 In vitro kinase assay and Kinase-Glo® luminescent kinase assay

GST-tagged CREB was expressed and purified from *E. coli* BL-21 (Invitrogen). FLAG-tagged CAMKV wildtype and mutants were stably expressed in SK-N-AS cell lines or overexpressed in HEK293T cells. FLAG-tagged CAMKV proteins were purified from cells lysed with RIPA lysis buffer using anti-FLAG antibody and protein A agarose beads, followed by three times washing with PBS and one time with kinase assay buffer (20 mM Tris/HCL, pH 7.5, 10 mM MnCl_2_, 10 mM MgCl_2_, 50 mM NaCl_2_ 1 mM dithiothreitol, 20 mM β-glycerophosphate, 1 mM NaF). The beads carrying immune complexes were resuspended in 50 μL of kinase assay buffer containing 100 mM ATP. GST-CREB was added to the mixtures, the reaction mixtures were incubated at 30°C for 30 minutes, and 4 × loading buffer was added. Proteins were resolved by SDS-PAGE and detected by western blot using an anti-phospho-CREB antibody. For Kinase-Glo® luminescent kinase assay, the kinase reaction mixtures were collected after incubation and added into equal volume Kinase-Glo® regent (V6711, Promega) for 10 minutes at room temperature, followed by the luminescence recording.

### 2.10 CCK-8 cell proliferation assay

Cell proliferation was determined using a Cell Counting Kit-8 (Dojindo Molecular Technologies) as described previously (*37*). In brief, cells were plated into 96-well plates at the concentration of 1×10^3^ cells per well, then incubated at 37°C for different time periods. Relative cell proliferation was quantified by adding 10 μL of Cell Counting Kit-8 solution. The absorbance was measured at 450 nm after 3-hour incubation at 37°C.

### 2.11 Crystal violet staining

Cells in 6-well plates were washed twice with ice-cold PBS, followed by fixation with ice-cold methanol for 10 minutes. Methanol was removed, and the plate was incubated with 0.5% crystal violet solution (made with 25% methanol) for 10 minutes at room temperature. The plate was rinsed with double distilled water and dried at room temperature. The stained colonies were photographed by using a microscope (Olympus).

### 2.12 3D culture assay

NB cells were resuspended in a 1:1 mixture of Matrigel (# CB-40230, Thermo Fisher Scientific) and cell culture medium, then plated in 24-well plates. After 8 days of culture, colonies were visualized, and the sizes of the colonies were measured under a microscope (Olympus).

### 2.13 Quantitative PCR analysis

Total RNA from NB cells was extracted using Trizol reagent (Life Technologies) and treated with RNase-free DNase (Roche) according to the manufacturer’s instructions. One μg of total RNA was converted to cDNA, and quantitative RT-PCR analyses were performed using SensiFAST™ SYBR® kit (Bioline) on an Applied Biosystems StepOne Plus instrument. The primers used are listed in table S4. Relative levels of gene expression were analyzed using the 2^-ΔΔCt^ method.

### 2.14 In situ proximity ligation assay (PLA)

The interaction between CAMKV and CREB was assessed in SK-N-AS cells using an in situ PLA assay (Duolink *in situ* red starter kit, Sigma-Aldrich) according to the manufacturer’s instructions. Briefly, SK-N-AS cells were fixed in 4% paraformaldehyde for 10 min at room temperature, followed by permeabilization with 0.5% Triton X-100. Cells were then blocked at 37°C for 30 minutes and incubated with primary antibodies against CAMKV (1:500) (rabbit), CREB (1:500) (mouse), or control mouse IgG (1:500), followed by washing and incubation with the secondary anti-mouse or anti-rabbit antibodies conjugated to PLA probes. After ligation and amplification, the red fluorescence indicating the interaction between CAMKV and CREB was visualized under a fluorescence microscope (Nikon).

### 2.15 Chromatin immunoprecipitation (ChIP)

ChIP assays were performed by using a Chromatin Immunoprecipitation (ChIP) kit (Active Motif). The MYCN-amplified cell line NGP and MYCN-non-amplified cell line SK-N-AS were cross-linked with 1% formaldehyde. Cells were then washed with ice-cold PBS, lysed, and sonicated on ice to produce sheared soluble chromatin. The soluble chromatin was precleared with Protein A Plus agarose beads (sc-2001, Santa Cruz Biotechnology) at 4°C for one hour and then incubated with anti-MYCN or anti-c-MYC antibodies or with control mouse IgG at 4°C overnight. The immunoprecipitated complexes were collected on protein A agarose and eluted. Crosslinking of immunoprecipitated chromatin complexes and input controls were reversed by heating at 65°C for 4 hours, followed by proteinase K treatment. The purified DNA was analyzed by quantitative PCR.

### 2.16 Immunohistochemistry (IHC)

IHC was performed on 30 formalin-fixed, paraffin-embedded human NB tissue specimens, including 19 high-risk NBs (11 with MYCN amplification and 8 without MYCN amplification) and 11 low-risk NBs. Sections were cut at 5 μm and put on the slides. The slides were then incubated at 70°C for 30 minutes. The sections on the slides were de-waxed into water, soaked in Leica Epitope Retrieval 1 (pH 6), and heated in pressure cook at 125 °C for 90 seconds (BioCare Medical), and then transferred into Leica Bond Rx Autostainer to run IHC staining using CAMKV monoclonal antibody 1:100 (#sc-102406, Santa Cruz) for 30 minutes. This study was approved by the Institutional Review Board of Children’s Hospital Los Angeles.

### 2.17 *In vivo* studies

NSG mice were obtained from The Jackson Laboratory and maintained at the Children’s National Hospital animal care facility. All treatments described were approved by the Institutional Animal Care and Use Committee (IACUC) at Children’s National Hospital under protocol number #000031057. To test the effect of *CAMKV* knockout on NB tumor growth, mice were injected intraperitoneally (i.p.) with 5 × 10^6^ firefly luciferase-labeled CHLA-136 (CHLA-136-Fluc) cells expressing the inducible *CAMKV* knockout plasmid or the control plasmid. Doxycycline was administered via intraperitoneal injection at 5 µg/mouse daily for three days. For in vivo therapeutic experiments, mice were injected i.p. with 5 × 10^6^ CHLA-136-Fluc NB cells. Two weeks after injection, mice were treated with i.p. injected OTSSP167 (10 mg/kg) daily for two weeks. Tumor growth was monitored weekly by bioluminescent imaging using a Xenogen IVIS Lumina system.

### 2.18 Mass spectrometry

#### Global proteome analysis

NGP wild-type and *CAMKV* knockdown cells were grown in RPMI-1640 with 20% fetal bovine serum, collected by centrifugation and washed with cold PBS and lysed with sonication using 10 sample volumes of 50 mM ammonium bicarbonate with 1 mM CaCl_2_. The protein concentration was determined using the Bradford assay. About 200 µg of lysate were digested with 4 µg of trypsin for 12 hours at 37°C. The concentration of tryptic peptides was measured using the Pierce™ Quantitative Peptide Assays kit (Thermo Fisher Scientific cat# 23275), and 25 µg of peptides were separated on a home-made reverse-phase C18 column in a pipet tip as previously (*38*). Peptides were eluted and separated into fifteen fractions using a stepwise gradient of increasing acetonitrile (2%, 4%, 6%, 8%, 10%, 12%, 14%, 16%, 18%, 20%, 22%, 24%, 26%, 28%, 35% acetonitrile) at pH 10. Subsequently, these fractions were combined into five groups (2%+12%+22%, 4%+14%+24%, 6%+16%+26%, 8%+18%+28%, 10%+20%+35% acetonitrile eluted fractions) and vacuum dried. The dried peptides were resuspended in a solution of 5% methanol and 0.1% formic acid in water, then subjected to a 2 cm trap column (100 µm i.d.) and separated by a 5 cm analytical column (150 µm i.d.) containing Reprosil-Pur Basic C18 (1.9 µm, Dr. Maisch GmbH, Germany). A nanoLC-1000 (Thermo Scientific, USA) delivered a 75 min discontinuous gradient of 4 to 24 % of acetonitrile/0.1% formic acid at a flow rate of 800 nL/min. An Orbitrap Fusion mass spectrometer (Thermo Scientific) was operated in data-dependent acquisition mode with the following parameters: MS was in the 300-1400 *m*/*z* range with a 120,000 resolution at 400 *m*/*z* and AGC target of 7.5 × 10^5^ (50 ms maximum injection time). Cycle time was top 3 selected MS1 signal using Quadrupole filter in 2 *m*/*z* isolation window, 2 seconds exclusion time. The HCD fragmented ions were detected by ion trap with rapid scan, 3 × 10^4^ AGC target, and 35 ms of maximum injection time.

#### Global phospho-proteome analysis

After trypsin digestion, 100 µg of peptides went through a phosphopeptide enrichment procedure Briefly, the peptide mixture was resuspended in 100 μL of binding buffer (50% acetonitrile, 1% trifluoroacetic acid, and 1M lactic acid) and incubated with 5 mg of pre-conditioned TiO2 beads (GL Science, 5020-75000) at room temperature for 20 minutes with continuous vortexing at 1,500 rpm, then spun at 500 × *g* for 30 seconds. The beads were washed with the binding buffer and collected by centrifuge at 500 × *g* for 30 seconds. The beads were further washed with a washing buffer (30% acetonitrile, 0.07% trifluoroacetic acid) twice, and transferred to a 200 μL pipet tip with Empore™ C18 disk (3M, 2215) plug. The beads were settled in the tip. Another 100 μL of wash buffer was added and washed, followed by elution with 150 μL of elution buffer (40% acetonitrile + 10% NH_4_OH). Then eluted samples were subjected to mass spectrometry analysis. The nanoLC-MS/MS and data analysis was carried out in the same condition as global profiling.

#### CAMKV phosphorylation site identification

Two µg of purified CAMKV protein was digested by 100 ng of trypsin (1:20 ratio) in the digestion buffer of 50 mM NH_4_HCO_3_, 1mM CaCl_2_ for 4 hours. Digested peptides were acidified by adding 2% formic acid (final concentration 0.2%) and loaded onto Orbitrap Fusion mass spectrometer. A targeted method (PRM, Parallel-Reaction Monitoring) was adopted to detect the defined peptides, including phosphorylated (EPCGpTPEYLAPEVVGR, 898.8999 m/z, charge 2+) and non-phosphorylated forms (EPCGTPEYLAPEVVGR, 858.9167 m/z, charge 2+). The peptides were isolated with a 45 min discontinuous gradient of 10-26% acetonitrile/0.1% formic acid at a flow rate of 850 nL/min, and directly introduced into the mass spectrometer. The MS was operated in data-dependent mode, the top 35 strongest ions were selected and fragmented under direct control of Xcalibur software (Thermo Scientific). Parent MS spectra were acquired in the Orbitrap with a resolution of 120,000, and selected ions were fragmented by HCD in the ion trap.

#### MS data processing

Raw MS data were processed by the Max-Quant software program (version 1.6.17.0; Max Planck Institute of Biochemistry, Martinsreid, Germany). Spectra were searched against target-decoy uniprot human reference proteome database (downloaded in July 2023 with a total of 20,408 Swiss-Prot reviewed canonical proteins) with a 1% of false-discovery rate at both protein and peptide levels. Carbamidomethyl(C) was selected as a fixed modification and Oxidation (M) and Acetyl (Protein N-term) were considered as variable modifications in global proteome analysis. A minimum peptide length was 7 amino acids. The MS1 mass tolerance was 20 ppm and the MS2 mass tolerance was 0.5 Dalton. A maximum of two missed cleavages was allowed. All other parameters were set to the default values. For global phospho-proteomic analysis, additional variable modifications (Phosphorylation of Ser, Thr and Tyr) were considered and a minimum site localization probability of 0.75 was required for localization of phosphorylation sites. MaxQuant LFQ algorithm for label-free quantification was used for peptides and proteins quantification. Peptide intensities was used for phosphopeptide quantification. Targeted phosphopeptides were validated by Skyline software and peak area (AUC) was calculated.

### 2.19 RNA sequencing

Total RNA was isolated by RNeasy Plus Mini Kit (Qiagen) from *CAMKV*-knockdown, OTSSP167 treated, or control cells. Quality was assessed before the RNA-seq library construction by Bioanalyzer RNA 6000 (Agilent Technologies, Inc). The library was sequenced on an Illumina HiSeq platform. Adaptors were removed from the raw reads, and sequencing quality was assessed with FastQC software (version 0.11.2). Quality scores, sequence duplication, adaptor content, and other metrics were examined to determine whether additional filtering was needed before the genome mapping. Clean reads were mapped onto the reference genome (GRCh38.p13) with the HISAT2 alignment program (version 2.1.0). The mappable reads were assembled into transcripts or genes with the StringTie assembler (version 1.3.5). Only coding genes were retained and considered for downstream analysis.

### 2.20 Bioinformatic and Statistical Analyses

To identify differentially expressed genes (DEGs) between *CAMKV*-knockdown and control cells or OTSSP167 treated vs untreated cells, we conducted DESeq2 (version 1.28.1) analyses in R software. Fold changes were log2-transformed, and adjusted P values (padj) were calculated using the Benjamini-Hochberg procedure to control the false discovery rate. Compared to the control cells, genes in the *CAMKV*-knockdown or OTSSP167 treated cells were considered significantly upregulated if the log2 fold-change was ≥ 1 with a padj < 0.05 or significantly downregulated if the log2 fold-change was ≤ –1 with a padj was < 0.05. Differentially expressed genes were selected for further analysis. For pathway analysis, Reactome and KEGG databases were chosen and DEGs were subjected to over-representation tests. DEGs were categorized into GO Biological Processes.

A student’s t-test was performed to identify differentially expressed proteins (DEPs) between CAMKV-knockdown and control cells. DEPs were called if the fold-change was ≥ 2 or <= 0.5 with a *p*<0.05. Reactome and KEGG databases were considered for pathway analysis, and DEPs were subjected to over-representation tests with all identified proteins as background.

To identify differentially expressed phosphosites between *CAMKV*-knockdown and control cells, DESeq2 (version 1.28.1) analysis using peptide intensities was conducted. Phophosites were considered significantly upregulated if the log2 fold-change was ≥ 1 with a padj < 0.05 or significantly downregulated if the log2 fold-change was ≤ –1 with a padj was < 0.05. Differentially expressed phosphosites were categorized into different protein classes (Function and compartment based) and kinase families based on the Human Protein Atlas Database (https://v19.proteinatlas.org/humanproteome/proteinclasses).

### 2.21 System Preparation and Molecular Docking Simulations

The structure of CAMKV was predicted using ColabFold (AlphaFold 2 colab) (*39*). The MELK/8a complex (PDB entry: 5IH9) was retrieved from the RSCB Protein Data Bank (*40*). Both protein structures were prepared using *Protein Preparation Wizard* via *Mastro* in Schrödinger (Schrödinger, LLC, New York, NY). Briefly, the preparation protocol included removing crystallographic water molecules, adding hydrogen atoms to the structure, and filling the missing side chains and loops. Then by using *Receptor Grid Generation*, a grid with size 10 × 10 × 10 Å was created within the MELK/8a complex to define the docking space with the bound ligand (8a) as the centroid following the Glide Grid Generation panel. For CAMKV, both Cys52 and Lys53 were selected as the centroid for generating a grid with the size of 25 × 25 × 25 Å.

For the ligands, MELK-8a (Compound CID: 119058124), K252a (Compound CID: 3813), and OTSSP167 (Compound CID: 135398499) were retrieved from the NIH PubChem. All ligands were subsequently prepared through the *LigPrep* module with the OPLS3 force field, the ionization state of inhibitors was assigned by Epik, pH was set to neutral 7.0 ± 2.0, and other configurations were set as default.

Molecular docking simulation was performed using the *Ligand Docking (Glide)* module via *maestro* in Schrödinger. The precision was set as XP (extra precision), the number of best poses per ligand keeping for energy minimization was set as 32, the number of poses per ligand to include to perform post-docking minimization was set as 16, and other configurations as default.

## 3. Results

### 3.1 CAMKV is required for CREB phosphorylation and NB cell proliferation

Since CREB has been demonstrated to upregulate multiple cell cycle-regulatory cyclin genes and promote cell growth, we investigated its role in NB cells. Examining CREB expression and its phosphorylation status in nine NB cell lines by immunoblotting assay revealed phosphorylation at Serine 133 in all cell lines (Fig.1A). CREB Knockdown using short hairpin RNA (shRNA) led to significant inhibition of cell proliferation in four NB cell lines (Fig.1, B and C, Fig.S1A). Further assessments of the CREB knockdown effects on cyclin gene expression showed a highest reduction of CCNA2 expression compared other cyclin genes in NB cells (Fig.1B, Fig.S1B). Additionally, CCNA2 knockdown in different NB cell lines also resulted in significant inhibition of NB cell proliferation (Fig.S1, C and D). These data highlight that CREB up-regulates CCNA2 expression and is critical for NB cell proliferation.

**Fig. 1.**
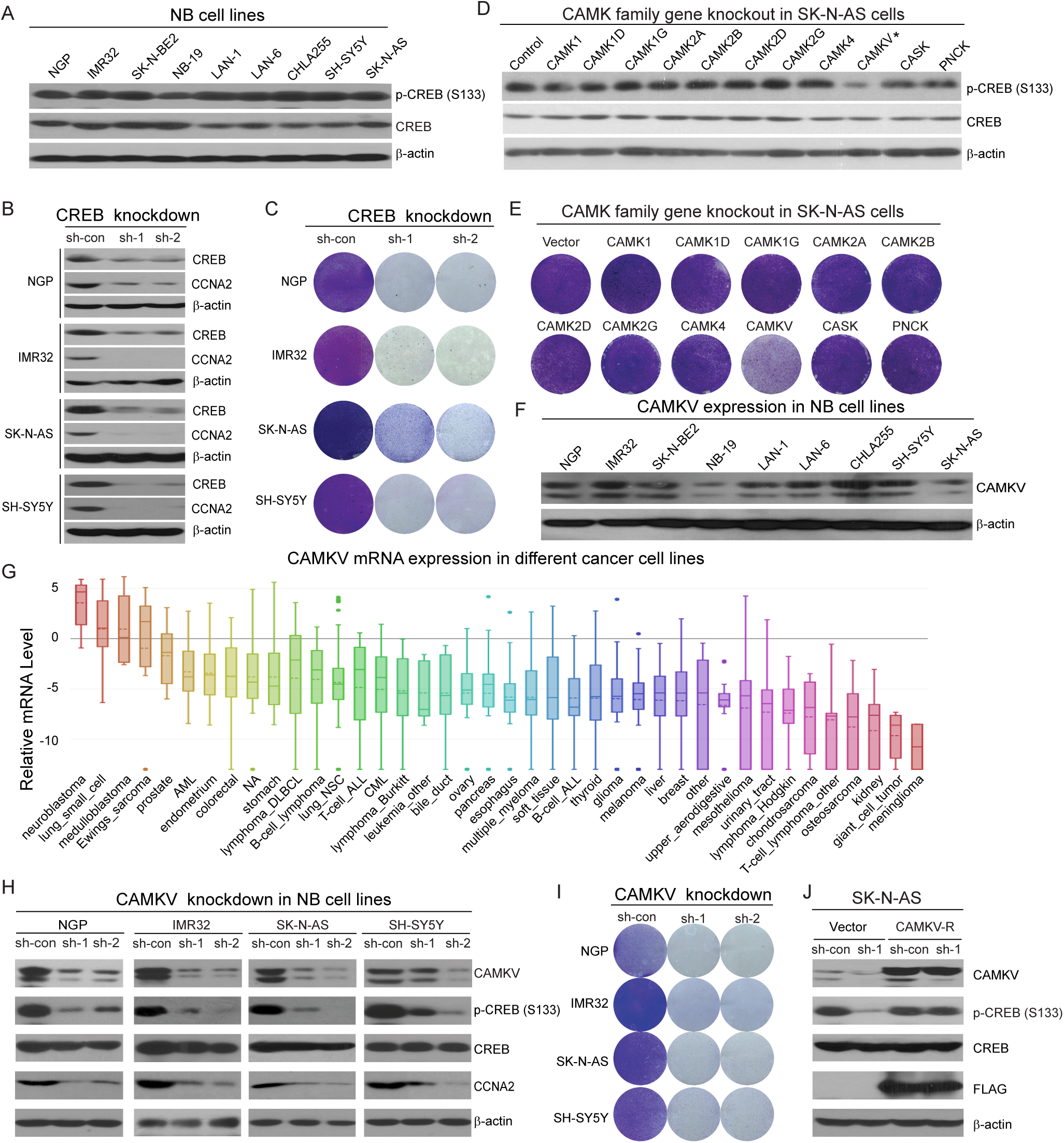
CAMKV is required for CREB phosphorylation and NB cell proliferation. **(A)** CREB phosphorylation level in NB cell lines. Immunoblot of cell extracts of nine NB cell lines to determine the level of total and phosphorylated-Ser133 CREB. β-actin was used as a loading control. **(B)** The effects of CREB knockdown by shRNA on CREB and CCNA2 expression. **(C)** The effect of CREB knockdown by two shRNA on NB cell proliferation by crystal violet staining after 7 days. **(D, E)** The effects of CRISPR/Cas9 knockout of each member of Ca^2+^/calmodulin-dependent kinases (CaMKs) on CREB phosphorylation (D) and cell proliferation (E) of SK-N-AS cells. **(F**) CAMKV protein levels in nine NB cell lines. **(G)** *CAMKV* mRNA expression in 40 tumor types in the Cancer Cell Line Encyclopedia (CCLE). **(H, I)** The effects of CAMKV knockdown on total and phosphorylated CREB and CCNA2 protein (H), as well as on the cell proliferation of NB cell lines by crystal violet staining assay (I). **(J)** The effects of CAMKV knockdown on CREB phosphorylation can be prevented by expressing a sh-RNA-resistant *CAMKV* (*CAMKV-R*) in SK-N-AS cells.

Several CaMK family members have been reported as CREB kinases in various tumor types andimplicated in cancer development (*41–44*). To explore whether the CaMK family of genes plays a role in NB cell proliferation, we individually knocked out the eleven CaMK family members using CRISPR-Cas9-mediated genome editing in SK-N-AS cells (Fig.1, D and E, Fig.S2). Of these, CAMKV knockout showed the most significant decrease in CREB phosphorylation and cell proliferation. CAMKV knockout was confirmed by genomic sequencing (Fig.S3).

Further, we detected CAMKV protein in nine tested NB cell lines, regardless of MYCN-amplification status (Fig.1F). Analyses of *CAMKV* mRNA expression from the Cancer Cell Line Encyclopedia (CCLE) of 40 tumor types revealed that NB has the highest CAMKV expression (Fig.1G). Consistent with the *CAMKV* knockout results, *CAMKV* knockdown led to decreased CREB phosphorylation and reduced CCNA2 levels (Fig.1H), while total CREB was unaffected by *CAMKV* knockdown. *CAMKV* knockdown also impaired cell proliferation in NB cell lines (Fig.1I). Importantly, expressing a shRNA-resistant *CAMKV* (CAMKV-R) in NB cells rescued the effect of *CAMKV* knockdown on CREB phosphorylation (Fig.1J). To confirm this knockdown result, we transfected piggyBac (PB) vector with an inducible CAMKV shRNA cassette and established stable line in SK-N-AS cells. Induction of CAMKV shRNA expression led to decreased CREB phosphorylation and cell proliferation (Fig.S4). Taken together, these results demonstrate that CAMKV is overexpressed in NB cells and plays a crucial role in CREB phosphorylation and cellular proliferation.

### 3.2 CAMKV is an active kinase in NB cells to promote cell proliferation

Our CAMKV immunoblots demonstrated two distinct bands in NB cell lines (Fig.1F), indicating the presence of two CAMKV isoforms. By sequencing CAMKV cDNA clones generated from NB cell lines, we identified the full-length isoform (CAMKV-FL) coding for 501 amino acids and a truncated isoform (CAMKV-S) with a 31-amino-acid C-terminal deletion (Fig.2A).

**Fig. 2.**
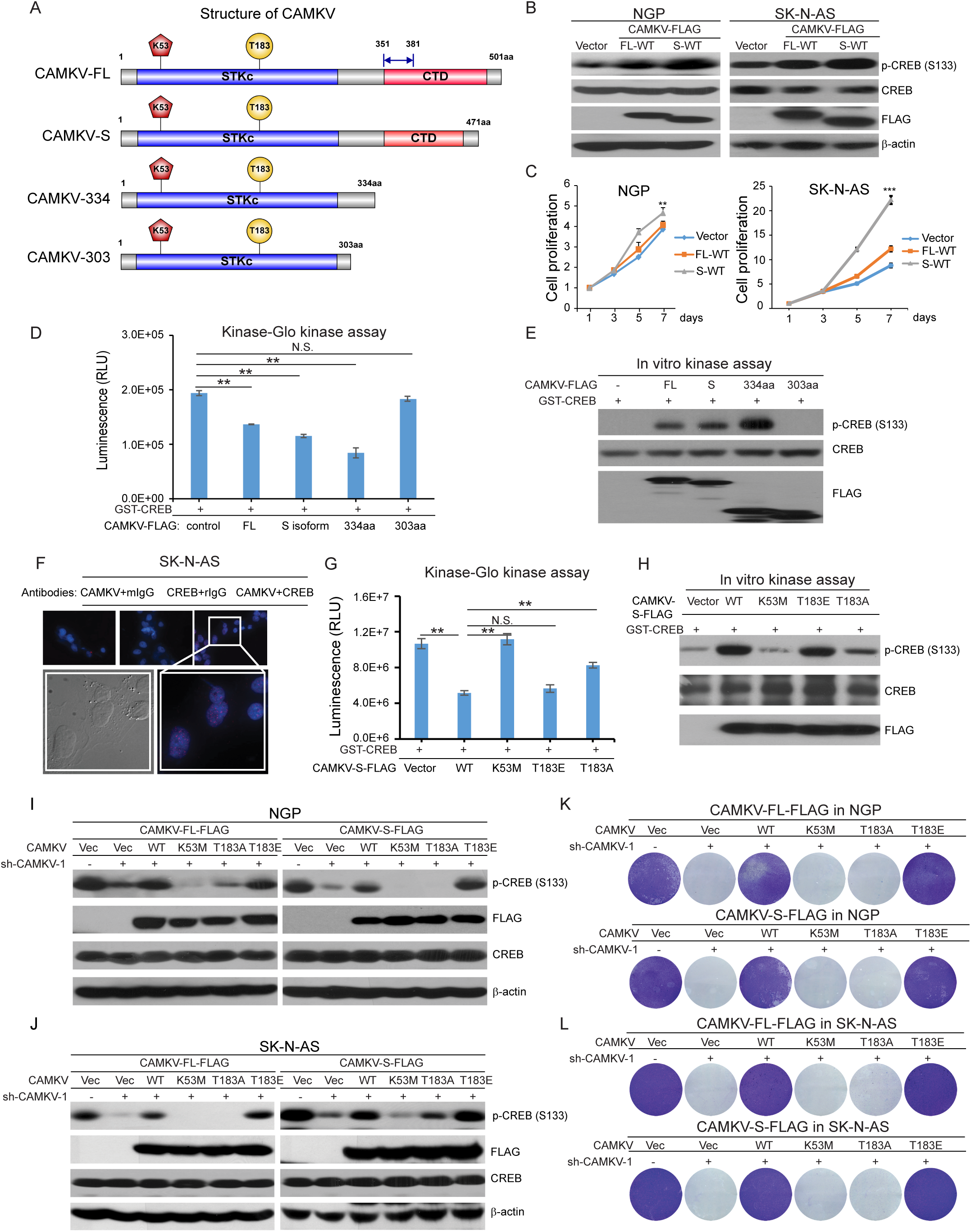
Phosphorylation at Thr183 is required for CAMKV kinase activation and signaling. **(A)** Diagram of CAMKV isoforms and truncated forms. The kinase domain (SPKc in blue) and regulatory domain (in red) containing CTD are illustrated. The missing amino acids (aa 351-381) in CAMKV short isoform (CAMKV-S) are indicated. The structures of truncated forms of CAMKV, CAMKV-334 and CAMKV-303 are also indicated. **(B, C)** The effects of over-expression of CAMKV-FL-WT or CAMKV-S-WT on CREB phosphorylation (B) and cell proliferation (C) of NGP and SK-N-AS cells. **(D, E)** In vitro phosphorylation of recombinant GST-CREB by CAMKV-FL, CAMKV-S, CAMKV-334aa, and CAMKV-303aa. CAMKV proteins were immunoprecipitated from cell lysates of SK-N-AS cells stably expressing CAMKV and then subjected to a Kinase-Glo® luminescent kinase assay (D) and an in vitro kinase assay (E). **(F)** Detection and visualization of the interaction between CAMKV and CREB in SK-N-AS cells by an in situ proximity ligation assay. **(G, H)** In vitro phosphorylation of recombinant GST-CREB by CAMKV-S-WT and putative kinase-dead K53M mutants, putative constitutively active kinase T183E mutants, and putative kinase-deficient T183A mutants in a Kinase-Glo® luminescent kinase assay (G) and an in vitro kinase assay (H). **(I-L)** The effects of stable expression of shRNA resistant CAMKV-FL, CAMKV-S or their mutants on CREB phosphorylation (I, J) and cell proliferation (K, L) in SK-N-AS and NGP cell lines, in which endogenous CAMKV was knocked down.

Both CAMKV isoforms contain an N-terminal protein serine/threonine kinase domain and a C-terminal regulatory domain containing a C-terminal repeat domain (CTD) with a characteristic Nona-peptide repeat TPA motif (Fig.2A). To determine the functional difference between CAMKV-FL and CAMKV-S, we ectopically overexpressed them in NB cell lines. While overexpression of both isoforms enhanced CREB phosphorylation, CAMKV-S overexpression resulted in stronger CREB phosphorylation and increased cell proliferation (Fig.2, B and C, Fig.S5, A and B).

A previous report suggested that CAMKV lacks kinase activity (*30*). However, our study indicated that CAMKV is crucial for CREB phosphorylation in NB cells. These data prompted us to investigate whether CAMKV acts as a functional kinase mediating CREB phosphorylation in NB cells. To determine whether CAMKV could directly consume ATP and phosphorylate CREB, we conducted both Kinase-Glo luminescent kinase assays and in vitro kinase assays with recombinant GST-CREB as the substrate. These assays revealed that purified FLAG-tagged CAMKV-FL, -S, and CAMKV-334aa (containing the kinase and calmodulin-binding domains) from the transfected SK-N-AS cells consumed ATP and directly phosphorylated recombinant CREB at Ser133 (Fig.2, D and E). However, a shorter form of CAMKV-303aa (kinase domain only) failed to phosphorylate CREB. Moreover, a cell-based in situ proximity ligation assay designed to visualize the spatial proximity of interacting proteins demonstrated a direct interaction between endogenous CAMKV and CREB in SK-N-AS cells (Fig.2F). Our results suggest that CAMKV directly phosphorylates CREB at Ser133 and the calmodulin-binding domain is essential for CAMKV kinase activation in NB cells. We followed up these experiments by overexpressing the various constructs in HEK293T cells, and we observed only CAMKV-334aa phosphorylated CREB in both Kinase-Glo and *in vitro* kinase assays (Fig.S5, C and D). Therefore, CAMKV-334aa protein was purified from HEK293T cells for phospho-proteomic analysis, and only Thr183 (T183) phosphorylation was identified (Fig.S5, E and F), suggesting that Thr183 plays a critical role in CAMKV kinase activation, and the calmodulin-binding domains is essential for its kinase activation. However, CAMKV-FL and CAMKV-S purified from HEK293T failed to phosphorylate CREB, suggesting that the C-terminal regulatory domain has inhibitory function and the mechanism leading to CAMKV-FL and CAMKV-S activation only exists in NB cells.

Phosphorylation and dephosphorylation of key serine and threonine residues in the activation loop of kinases are crucial for regulating kinase activity (*45–47*). Numerous studies have demonstrated that substituting these critical residues with acidic residues to mimic phosphorylation renders the kinase constitutively active (*48*, *49*). To assess whether CAMKV kinase activation depends on the K53 (potential site for ATP binding) and T183 (potential phosphorylation site for kinase activation), we generated FLAG-tagged mutant expression constructs: putative kinase-dead K53M mutant, constitutively active kinase T183E mutant, and kinase-deficient T183A mutant. We found that CAMKV-S-WT and CAMKV-S-T183E proteins, purified from transfected SK-N-AS cells, directly phosphorylated recombinant CREB at Ser133, with lesser phosphorylation by CAMKV - S-T183A protein (Fig. 2, G and H). In contrast, CAMKV-S-K53M and CAMKV-S-T183A proteins failed to phosphorylate CREB (Fig. 2, G and H). Overexpression of CAMKV-WT and -T183E enhanced CREB phosphorylation and promoted NB cell proliferation compared to the control vector, while CAMKV-K53M and -T183A inhibited CREB phosphorylation and cell proliferation (Fig.S5, G and H). Moreover, ectopic expression of the shRNA-resistant CAMKV-WT or CAMKV-T183E rescued the defective CREB phosphorylation and cell proliferation caused by knocking down endogenous CAMKV in NB cells, whereas K53M and T183A mutants failed to do so (Fig.2, I to L).

Three-dimensional cell culture systems accurately model physiological cell growth and cell-cell interactions. SK-N-AS cells with CAMKV-FL-WT and CAMKV-S-WT overexpression formed larger colonies compared to the vector control cells, while cells with K53M mutant overexpression formed smaller colonies (Fig.S5, I and J). Notably, CAMKV-S-WT overexpression led to the formation of larger colonies than CAMKV-FL-WT, indicating that the CAMKV-S isoform is more active in promoting NB cell proliferation (Fig.S5, I and J). These results suggest that the deleted 31-amino acid sequence in the CAMKV-S isoform inhibits its kinase activity.

We also investigated the role of the calmodulin binding domain in CAMKV kinase activation. Alignment of the calmodulin binding domains from human CAMK family members revealed several conserved residues including three lysine and one arginine residues (Fig.S6A). Substituting them with alanine residues inactivated CAMKV kinase activity in NB cells (Fig.S6, B and C). However, this substitution had no effect on the constitutively active CAMKV (CAMKV-T183E-R3K-mut) (Fig.S6, B and C). Overexpressing shRNA-resistant CAMKV-T183E with the four sites substituted with alanine (T183E-R3K-mut) also rescued the defective CREB phosphorylation and cell proliferation caused by knocking down endogenous CAMKV in SK-N-AS cells (Fig.S6, D and E). In contrast, CAMKV-WT with the four sites substituted with alanine (WT-R3K-mut) did not rescue these effects (Fig.S6, D and E). These results indicate that the calmodulin-binding domain of CAMKV is required for the initial kinase activation, but not for maintaining kinase activity after the kinase phosphorylation and activation. Collectively, these findings suggest that at least a portion of the C-terminal regulatory domain plays an inhibitory role, and the calmodulin-binding domain is essential for CAMKV phosphorylation at T183 and its subsequent activation in NB cells.

Interestingly, an earlier study suggested that CAMK4 acts as a CREB kinase in the NB cell line, SK-N-BE2 (*50*). However, we found that overexpressing CAMK4 in SH-SY5Y, SK-N-AS, and NGP cell lines inhibited NB cell proliferation while having no significant effect on CREB phosphorylation in NB cells (Fig.S7, A and B). Furthermore, FLAG-CAMK4 purified from SK-N-AS cells failed to consume ATP and phosphorylate CREB in both the Kinase-Glo luminescent kinase assay and the in vitro kinase assay (Fig.S7, C and D). Additionally, CAMK4 gene knockout had no effect on CREB phosphorylation or cell proliferation in NB cells, suggesting that CAMK4 is not a CREB kinase in this context (Fig.S7, E and F). CAMK4 knockout was confirmed by genomic sequencing (Fig.S8).

### 3.3 *CAMKV* is a direct transcriptional target of MYCN/MYC in NB cells

Myc family members MYCN and MYC share sequence homology and bind to chromatin at consensus E-box sequences (CAGCTG) as heterodimers with the bHLH Zip protein Max (*51*, *52*). To investigate whether MYCN/MYC directly regulate *CAMKV* gene expressions, we analyzed a previously generated NB ChIP-sequencing dataset (*53*) and identified two potential MYCN/MYC binding sites in the 5’ untranslated region (UTR) and promoter region of *CAMKV* (Fig.3A). We further validated these findings by performing individual ChIP-qPCR assays using an anti-MycN antibody for NGP cells (*MYCN*-amplified) and an anti-Myc antibody for SK-N-AS cells (Myc overexpressed). We observed significant enrichment of MYCN/MYC binding at the *CAMKV* promoter (Fig.3B). Analyses of the NB RNA-seq dataset GSE19274 (*51*) revealed higher *CAMKV* mRNA levels in *MYCN*-amplified NB cell lines compared to *MYCN*-non-amplified lines (Fig.3C). A significant positive correlation between *CAMKV* and *MYCN* expression was also observed (Pearson correlation, R=0.523) (Fig.3D). Furthermore, immunoblotting assays revealed an overall positive correlation of CAMKV and phospho-CREB with MYCN/MYC protein levels in NB cell lines (Fig.3, E and F). Additionally, *MYCN* and *MYC* knockdown led to decreased *CAMKV* expression levels (Fig.3G). Furthermore, *CAMKV* mRNA levels were higher in tumors than in normal ganglia and adrenal glands from the TH-MYCN-driven NB mouse model (*54*, *55*) (Fig.3, H and I). Taken together, these data suggest that *CAMKV* is a direct transcriptional target of MYCN/MYC in NB.

**Fig. 3.**
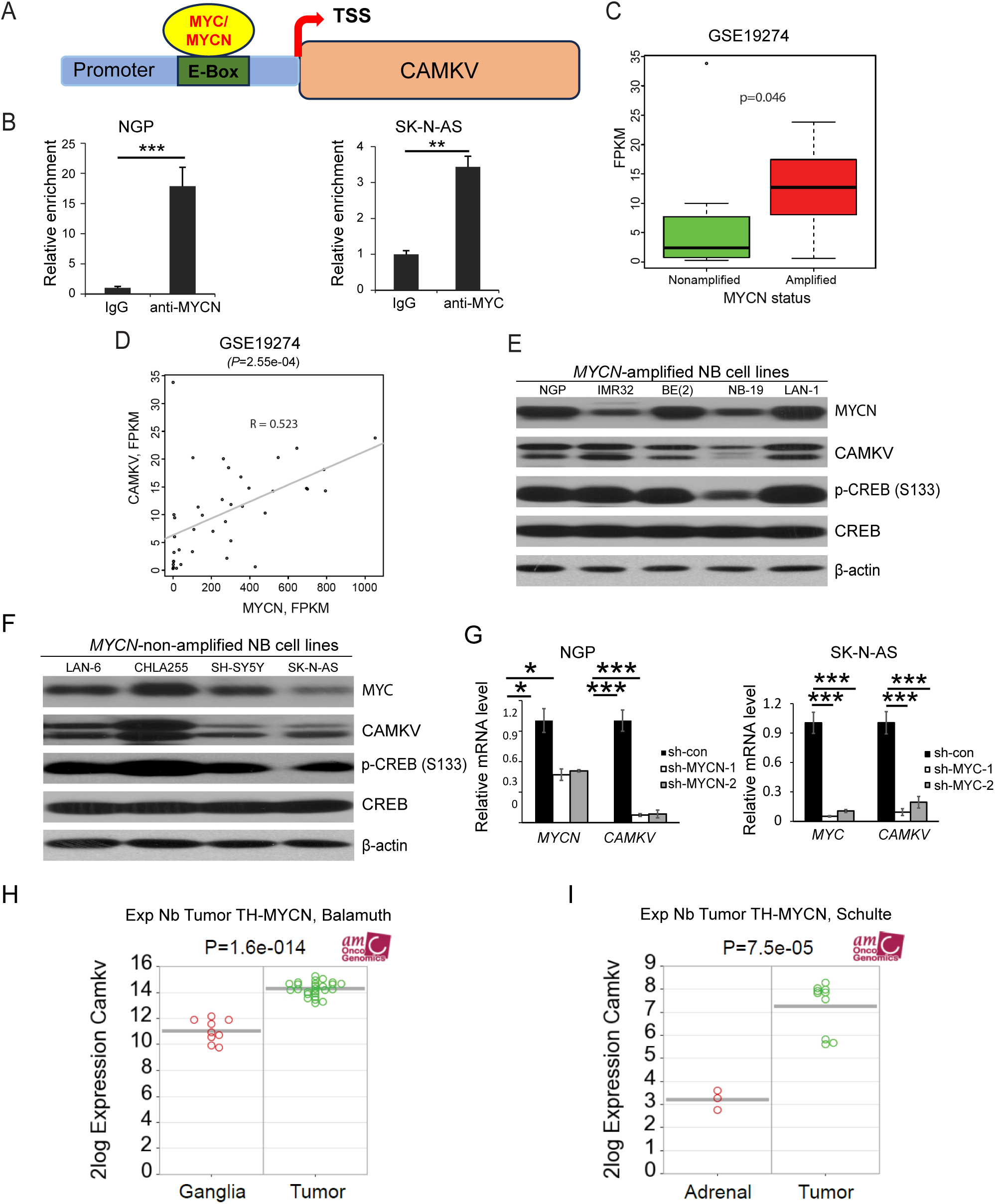
CAMKV is a direct transcriptional target of MYCN/MYC in NB cells. **(A)** Potential MYCN/MYC binding sites at the 5’UTR and promoter region of *CAMKV*. **(B)** ChIPs were performed with anti-MYCN and anti-MYC antibodies on two NB cell lines followed by quantitative PCR to detect the relative enrichments of target sequences. **(C)** The mRNA level of *CAMKV* in MYCN-amplified NB cell lines and MYCN-non-amplified NB cell lines (from GSE19274 dataset). **(D)** The correlation of gene expression between *CAMKV* and *MYCN* in NB cell lines (from GSE19274 dataset). **(E, F)** CAMKV and p-CREB protein levels in *MYCN*-amplified (E) and *MYCN*-non-amplified NB cell lines (F). **(G)** The effects of *MYCN/MYC* knockdown on *CAMKV* mRNA expression in NB cells. **(H-I)** The relative *Camkv* mRNA expressions in tumors derived from *TH-MYCN*-driven mouse model compared to ganglia and adrenal by analyzing Series GSE17740 (H) and GSE32386 (I) datasets.

### 3.4. *CAMKV* knockdown induces extensive transcriptomic and proteomic changes in NB cells

To elucidate how CAMKV regulates cell proliferation in NB and identify its downstream pathways, we conducted bulk RNA-sequencing (RNA-seq) on wild-type and *CAMKV* knockdown NGP cells (Fig.4A). Compared to wild-type cells, we identified 477 upregulated and 811 downregulated genes in *CAMKV* knockdown cells (table S5). Gene set enrichment analyses (GSEA) of downregulated genes revealed multiple linked neuronal pathways, suggesting CAMKV’s pivotal role in regulating the neuronal system (Fig.4B). Additionally, we found that many of the previously reported CREB target genes in table S6 were downregulated in *CAMKV* knockdown cells, confirming CREB as one of CAMKV’s targets (*56*). To identify CAMKV regulatory proteins and their associated pathways, we conducted a global proteome analysis that revealed 380 upregulated and 111 downregulated proteins in *CAMKV* knockdown cells compared to wild-type NGP cells (Fig.4C and table S7). GSEA of downregulated proteins identified multiple pathways related to protein translation, including translation initiation, elongation, and termination, as well as apoptosis (Fig.4D).

**Fig. 4.**
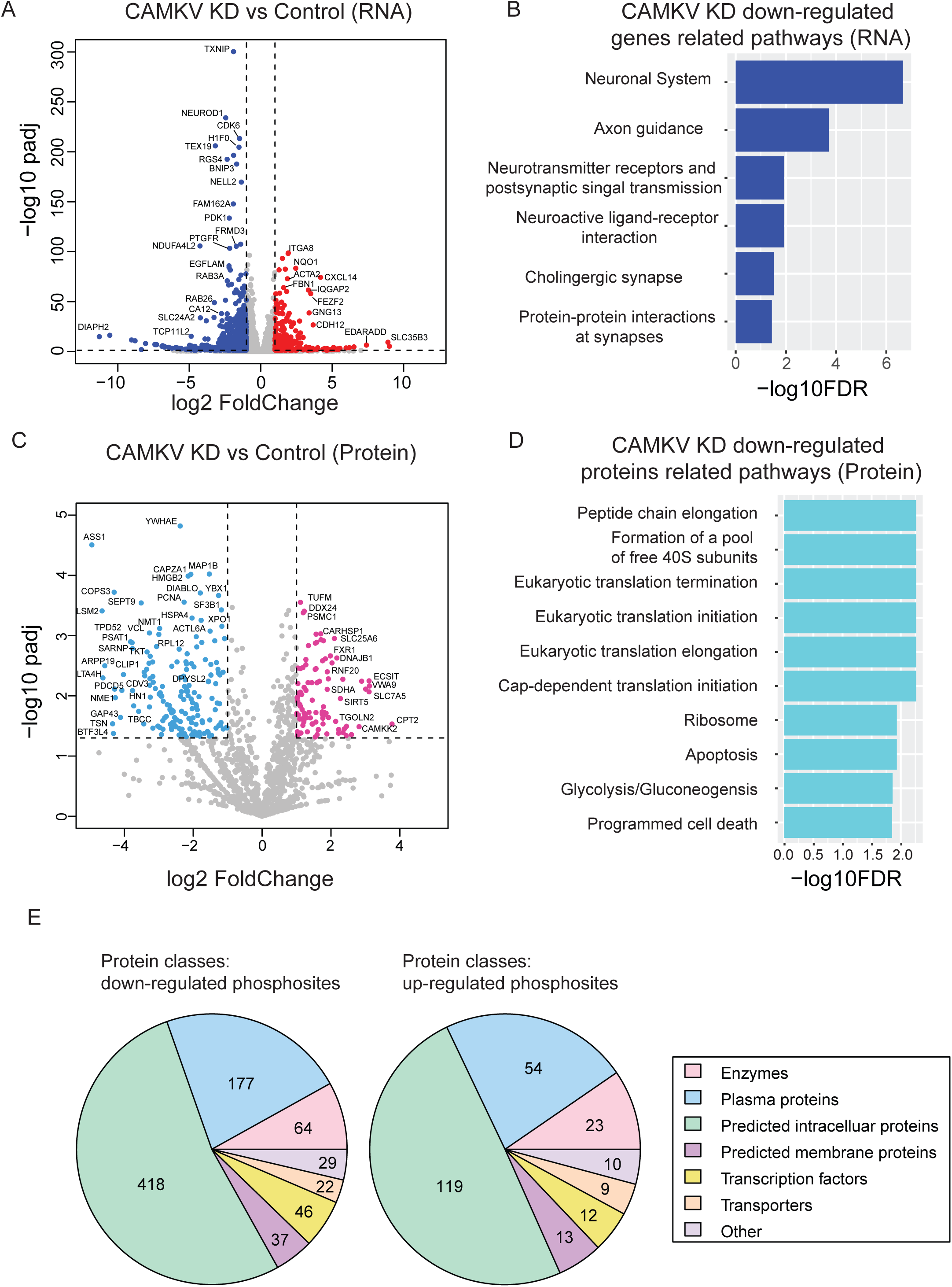
*CAMKV* knockdown induces extensive transcriptomic and proteomic changes in NB. **(A)** RNA-seq results of wild-type NGP cells and *CAMKV* knockdown cells. Plot of differentially expressed genes comparing significance vs fold change. **(B)** Gene set enrichment analysis of downregulated genes. **(C)** Global proteome analysis of wild-type and *CAMKV* knockdown NGP cells. Plot of differentially expressed proteins comparing significance vs fold change. **(D)** Gene set enrichment analysis of downregulated proteins. **(E)** Global quantitative phosphoproteomic analysis of WT and *CAMKV* knockdown NGP cells showing the protein classes with down- and up-regulated phosphosites.

Perturbing kinase activity in living cells through drug treatment or knocking down/out a specific kinase allows for examining consequent changes in phosphorylation levels in vitro (*57*, *58*). To systematically identify CAMKV substrates, we performed global quantitative phosphoproteomic analyses using NGP wild-type and *CAMKV* knockdown cells. These analyses revealed 240 upregulated and 793 downregulated phosphopeptides in *CAMKV* knockdown cells (Fig.4E and table S8). According to the Human Proteome Atlas database, these phosphosites fall into six protein classes: predicted intracellular proteins, plasma proteins, enzymes, predicted membrane proteins, transcription factors, and transporters. The enzymes include kinases, ubiquitin ligases and deubiquitinases, histone modifiers, DNA topoisomerases, RNA helicases, and enzymes involved in metabolism. These results suggest that CAMKV regulates NB cell proliferation via phosphorylating different proteins.

Noticeably, we identified 46 transcription factors with decreased phosphorylation changes following *CAMKV* knockdown (table S8 and S9). The decreased phosphorylation of transcription factor GATA2 at Ser182 and Ser192 in *CAMKV*-knockdown cells suggests that GATA2 is a potential substrate of CAMKV. Since anti-GATA2 phospho-Ser192 antibody is commercially available and GATA2 is highly expressed in NB cells, we validated this finding using immunoblotting assays (Fig.S9 and Fig.S10A). To further investigate the role of CAMKV’s kinase activity in GATA2 phosphorylation, we co-transfected FLAG-tagged GATA2 with CAMKV-334aa-WT or CAMKV-334aa-K53M into HEK-293T cells. We found that CAMKV-334aa-WT, but not CAMKV-334aa-K53M kinase-dead mutant, increased the phospho-Ser192-GATA2 level (Fig.S10B). In addition, GATA2 knockdown significantly inhibited cell proliferation compared to control cells (Fig.S10, C and D). Together, these findings suggest that GATA2 is a CAMKV kinase activity-regulated transcription factor and promotes NB cell proliferation.

### 3.5 *CAMKV* overexpression in NB is associated with poor patient survival

By analyzing a publicly available NB patient tumor dataset (https://pob.abcc.ncifcrf.gov/cgi-bin/JK), we found that *CAMKV* is highly expressed in both NB tumors and established cell lines, compared to other pediatric cancers (Fig.S11A). RNA-seq analysis of 498 NB patient tumor samples (https://hgserver1.amc.nl/) (GSE62564, Seqc-498-cohort) revealed a negative correlation between *CAMKV* gene expression and patient survival (p=1.0e-10) (Fig.5A). This finding was confirmed in another dataset of 649 patients (GSE45547, Kocak-649-cohort) (Fig.S11B). Additionally, we found that *CAMKV* mRNA levels were significantly higher in high-risk NB patients compared to low-risk patients (Fig.5B). Furthermore, *CAMKV* mRNA levels were significantly higher in tumors of advanced stages (stages 3, 4, and 4S) compared to those of low stages in both datasets (Fig.5C, Fig.S11C). We also observed that *CAMKV* mRNA levels were significantly higher in *MYCN*-amplified NBs compared to non-amplified ones (Fig.5D, Fig.S11D). In both datasets (GSE62564 and GSE45547), a positive correlation between *CAMKV* and *MYCN* mRNA levels was observed (R=0.494, p=4.7e-32 in the Seqc-498 cohort; R=0.471, p=3.4e-37 in the Kocak-649 cohort) (Fig.5E, Fig.S11E). In a single cell study of 16 NBs, 2 embryos, and 4 fetal adrenal glands (*59*), *CAMKV* was found to be highly expressed in neuroblasts (cell markers: *CHGA, TH, NCAM1*) from embryos, fetal adrenal glands (Fig.5F). *CAMKV* is overexpressed and positively correlated with *MYCN* amplification in NB cells identified by NB cell markers of *CHGA, TH, MYCN* (Fig.5F).

**Fig. 5.**
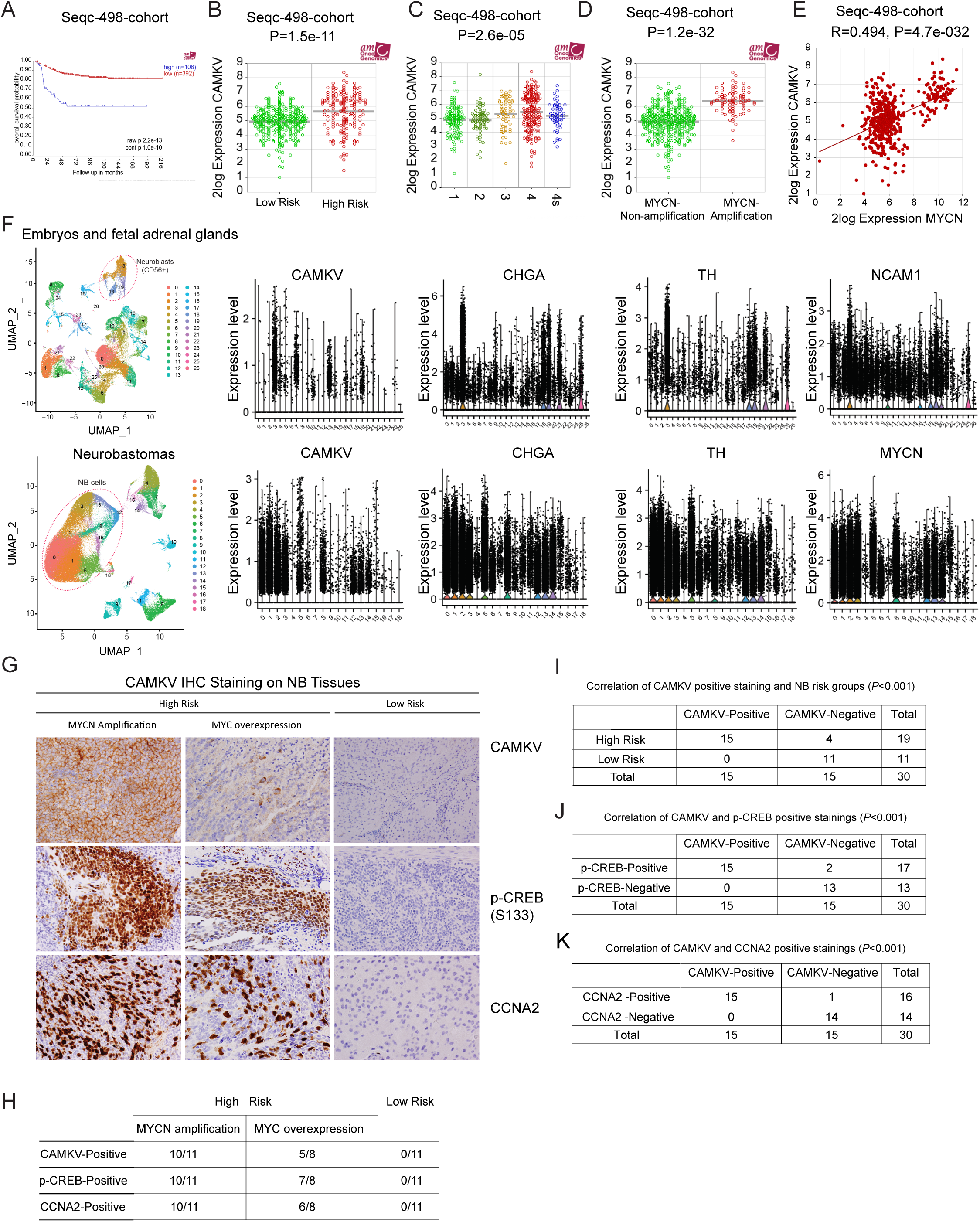
*CAMKV* high expression in NB tumors correlates with poor patient survival. **(A)** Correlation of high and low *CAMKV* mRNA levels of NB tumor samples and the overall survival of patients was analyzed using the GSE62564 (Seqc-498-cohort) dataset. **(B)** *CAMKV* expression in low- (n=322) and high-risk (n=176) NB samples from the Seqc-498-cohort. Statistical analysis was performed using the two-sided unpaired *t-*test. **(C)** *CAMKV* mRNA levels in patient tumor samples from the Seqc-498-cohort grouped on the base of INSS stage 1 to 4. Statistical analysis was performed using the two-sided unpaired *t* test. (Stage 1, n=121; Stage 2, n=78; Stage 3, n=63; Stage 4, n=183; Stage 4s, n=53). **(D)** *CAMKV* mRNA levels in tumors from patients with non-amplified *MYCN* (n=401) and amplified *MYCN* (n=92, GSE62564 dataset). **(E)** The correlation of gene expression between *CAMKV* and *MYCN* in the Seqc-498 cohort. **(F)** CAMKV and MYCN expressions in neuroblast and NB cells by single cell RNA-Seq of 2 embryos, 4 fetal adrenal glands, and 16 NBs, **(G)** Representative images of CAMKV, phospho-CREB, and CCNA2 IHC staining on the sections from 30 human NB patient tumor tissues. **(H)** Summary of IHC staining results. **(I-K)** The correlation between CAMKV protein expression and NB risk (I), the expression of p-CREB (S133) (J) and CCNA2 (K) from these tumors.

Consistently, IHC staining showed co-positive staining of CAMKV, p-CREB and CCNA2 in the high-risk human NB primary tumor samples (Fig.5, G and H). In 19 high-risk NB patients, CAMKV, p-CREB and CCNA2 staining were positive, not only in tumors with *MYCN* amplification, but also in tumors without *MYCN* amplification but with c-MYC overexpression (Fig.5H). In contrast, CAMKV, p-CREB and CCNA2 staining were negative in all low-risk tumors (Fig.5H). Based on these data, a strong correlation (*P*˂0.001) was observed between CAMKV positive staining and high-risk NB (Fig.5I), between CAMKV and p-CREB positive staining (Fig.5J), and between CAMKV and CCNA2 positive staining (Fig.5K).

In various tumor types, CAMK2A, CAMK2B, CAMK2D, CAMK2G, and CAMK4 have been reported to phosphorylate CREB (*28*, *29*). However, we found that high expression of these other CaMKs in NB primary tumor specimens predicted better patient outcomes (Fig.S12). The mRNA level of *CAMKV*, but not other CaMK family members, was significantly higher in *MYCN*- amplified NB patient tumor samples compared to *MYCN*-non-amplified ones (Fig.S13, A and B). Furthermore, the mRNA level of *CAMKV* was significantly higher in Stage 3 and Stage 4 NB patient tumors than in Stage 1 and Stage 2 samples (Fig.S13, C and D). Together, these data suggest that CAMKV potentially serves as a novel marker for risk stratification in NB prognosis.

### 3.6 Inhibition of CAMKV activity suppresses NB growth in vivo

The traditional CRISPR-Cas9 system lacks the ability to achieve temporally or spatially controlled Cas9 protein expression in vivo. Additionally, uncontrolled Cas9 expression in animal models can lead to genomic damage (*60*), off-target effects (*61*, *62*), and immunological clearance responses hindering the system’s application (*63*). To test the effect of CAMKV inhibition on NB growth in vivo, we employed a conditional CRISPR/Cas9-mediated gene knockout approach (*64–67*). This approach allows for temporal control of CRISPR-Cas9 activity for inducible genome editing in luciferase-expressing CHLA136 cells (CHLA136-Fluc). We found that doxycycline-regulated sgRNA and Cas9 induction facilitated efficient gene disruption in CHLA136-Fluc cells and led to decreased cell proliferation (Fig.6, A and B). We subsequently examined the effect of inducible *CAMKV* gene knockout on NB tumor growth in the CHLA136-Fluc xenograft model. Inducible *CAMKV* gene knockout resulted in decreased tumor growth (Fig.6C) and significantly prolonged the survival of mice carrying tumors (Fig.6D). These findings further demonstrate that CAMKV plays a significant role in NB growth.

**Fig. 6.**
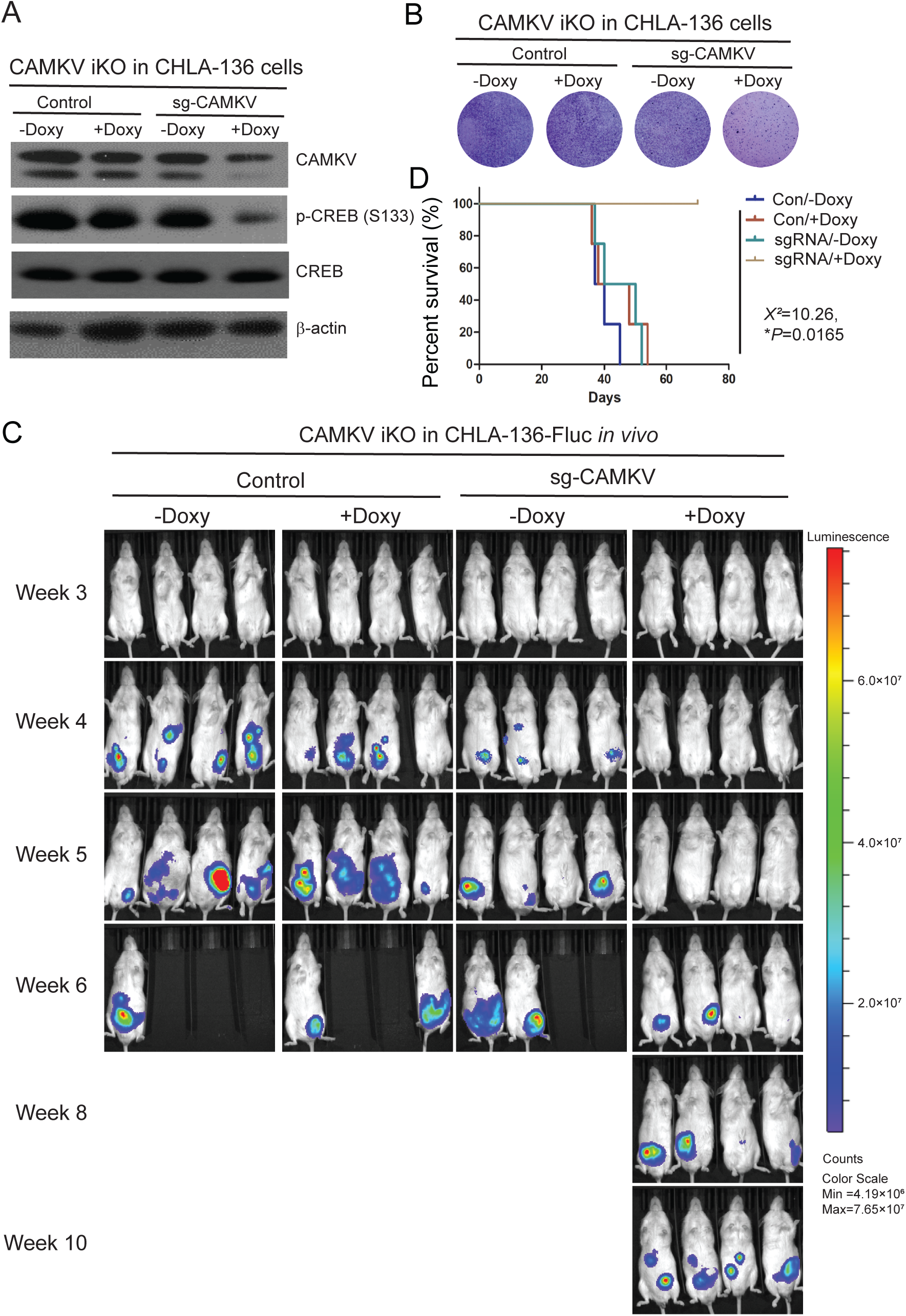
Inducible knockout of *CAMKV* gene suppresses NB cell proliferation in vitro and tumor growth in the xenograft NB mouse model. **(A)** Immunoblotting of CAMKV and p-CREB (S133) in CHLA-136 cell with inducible *CAMKV* gene knockout or control vector. **(B)** The effects of inducible knockout of *CAMKV* gene on NB cell proliferation. CHLA-136 cells were treated with doxycycline (100 ng/mL, 48 hours) and stained with crystal violet 14 days after treatment. **(C)** Bioluminescent imaging of CHLA-136-Fluc xenograft NB mice from the control group and the *CAMKV* inducible knockout group. **(D)** The survival rate of CHLA-136-Fluc xenograft NB mice of control group and the CAMKV inducible knockout group. Statistical analysis was performed by a Log-rank test. (χ^2^ = 10.26; p = 0.0165).

We serendipitously found that OTSSP167 and K252a inhibited ATP consumption by examining the inhibitory effect of 21 small molecules we have on the kinase activity of purified CAMKV-334aa in a Kinase-Glo assay (Fig.S14A). Our previous study has shown that OTSSP167 suppresses cell proliferation by inhibiting MELK activity in NB (*68*, *69*). OTSSP167 also inhibited cell proliferation in NB patient-derived xenograft (PDX) cells with a low IC50 (Fig.S14, B and C). In our CAMKV Kinase-Glo assay, OSSTP167 significantly suppressed ATP consumption and CREB phosphorylation in a dose-dependent manner (Fig.7, A and B). These results suggest that OSSTP167 is a CAMKV inhibitor in vitro.

**Fig. 7.**
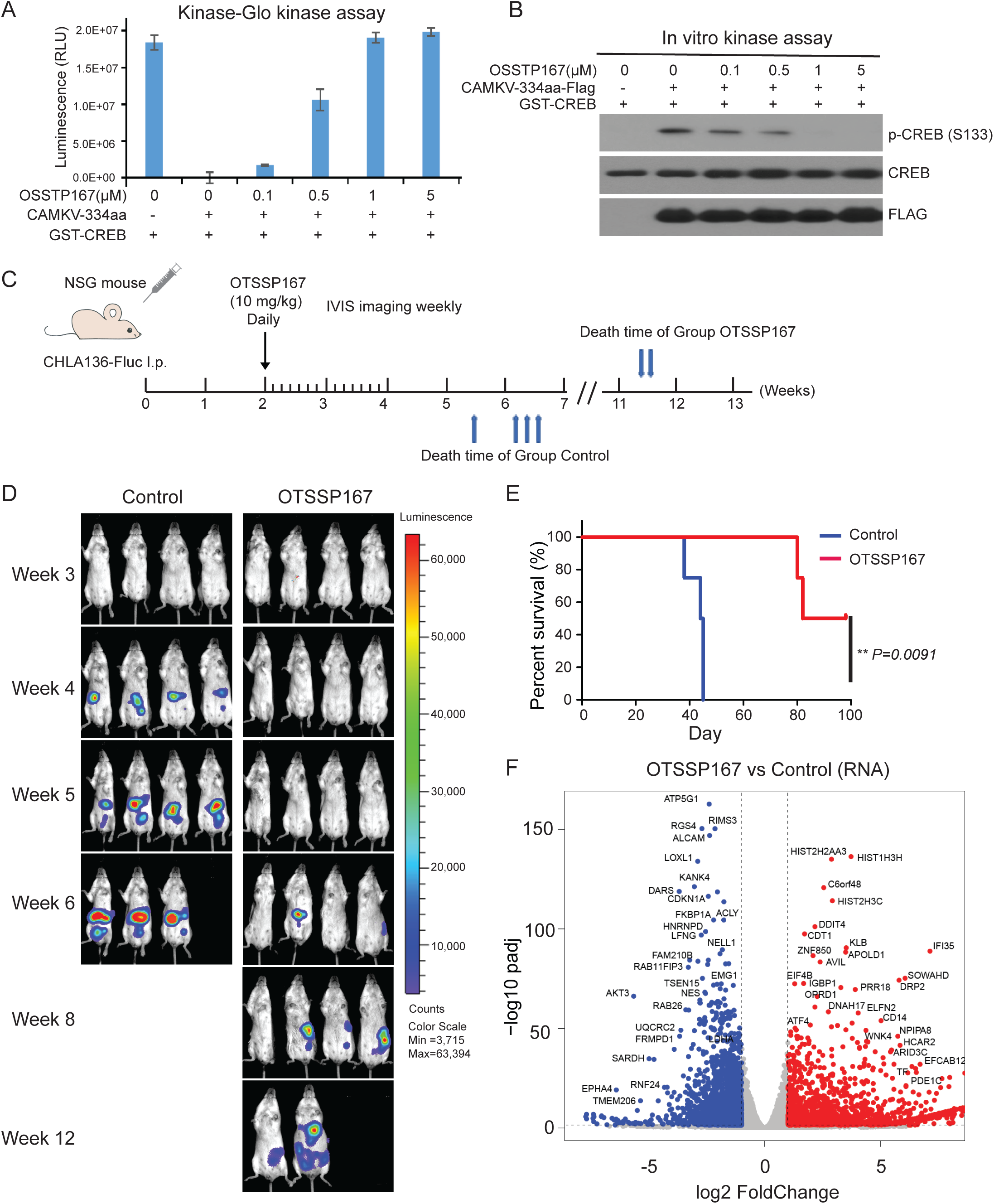
Inhibition of CAMKV activity by small molecular inhibitors suppresses NB cell proliferation in vitro and tumor growth in xenograft NB mouse model. **(A)** Kinase-Glo kinase assay to show the inhibition of OTSSP167 on CAMKV activity. **(B)** In vitro Kinase assay to show the inhibition of OTSSP167 on CREB phosphorylation mediated by CAMKV. **(C)** Treatment regimen of the xenograft NB mouse model. Two weeks after CHLA-136-Fluc cell i.p. implantation in NSG mice, OTSSP167 was i.p. administrated daily for 2 weeks. **(D)** Bioluminescent imaging photos of CHLA-136-Fluc xenograft NB mice from the control and the OTSSP167-treated groups. **(E)** The survival rate of CHLA-136-Fluc xenograft NB mice of control-(n=4) and the OTSSP167-treated groups (n=4). **(F)** RNA-seq results of control cells and OTSSP167-treated cells (60 nM, 12 hours). Plot of differentially expressed genes comparing significance vs fold change.

To evaluate the therapeutic potential of OTSSP167 in treating NB, we examined its activity in our CHLA136-Fluc xenograft model. Mice were randomized into different cohorts and treated with OSSTP167 (10 mg/kg) or vehicle daily for two weeks (Fig.7C). We found that OTSSP167 treatment significantly inhibited NB tumor growth and prolonged the survival of the mice compared to the vehicle-treated group (Fig.7, D and E).

To decipher the inhibitory mechanism of OTSSP167 in NB, we performed RNA-seq of NGP cells treated with OTSSP167 (60 nM, 12 hours). Our analysis revealed genome-wide transcriptome changes with 1,784 upregulated and 2,234 downregulated genes (Fig.7F and table S10). GSEA of the downregulated genes revealed pathways related to the neuronal system, like those observed in *CAMKV* knockdown cells (Fig.S14D). Gene Ontology (GO) term analysis of these genes revealed neuronal biological processes such as axon guidance, positive regulation of long-term synaptic potentiation, positive regulation of synaptic transmission, axonogenesis, and neuron projection guidance (Fig.S14E). Notably, 237 of the downregulated genes were also observed in *CAMKV* knockdown cells (Fig.S14F). Together, these data and our previously published data demonstrate the ability of OTSSP167 to inhibit NB tumor growth and prolong survival by dual targeting both CAMKV and MELK.

Several small molecule inhibitors have been identified for members of the CaMK family (*70–75*). We examined the inhibitory effects of K252a, K252c, KN93, STO609, PKC412 (midostaurin), and staurosporine on the kinase activity of purified CAMKV-334aa-FLAG-His using an in vitro kinase assay. Of these, only K252a and staurosporine inhibited CAMKV-334aa-mediated CREB phosphorylation in the in vitro kinase assay (Fig.S15A) and in the Kinase-Glo assay (Fig.S15B). Similarly, K252a also suppressed ATP consumption by the CAMKV short isoform (CAMKV-S-FLAG) kinase purified from SK-N-AS cells (Fig.S15C). Consistently, K252a and staurosporine, but not the others, inhibited cell proliferation (Fig.S15D), CREB phosphorylation (Fig.S15E), and CCNA2 expression (Fig.S15F), as well as reduced cell viability (Fig.S15G). These findings indicate that K252a suppresses NB cell proliferation at least partially by targeting CAMKV signaling.

Further, we performed molecular docking of OTSSP167, K252a and another MELK inhibitor 8a (*76*) and aligned the protein structure of CAMKV, especially at the ATP-binding pocket. The docking poses of OTSSP167, 8a and K252a within the ATP-binding pocket of CAMKV predict that they are strong inhibitors of CAMKV, and comparatively OTSSP167 (Docking Scores: −4.445 kcal/mol) had a highest docking score compared to 8a (Docking Scores: −3.543 kcal/mol) and K252a (Docking Scores: −2.640 kcal/mol) (Fig.S16). Together, these results suggest that inhibiting CAMKV kinase activity represents a promising therapeutic strategy for NB treatment.

## 4. Discussion

This study has identified CAMKV as a protein kinase and demonstrated its direct phosphorylation of CREB at S133, thereby promoting NB cell proliferation and tumor growth. Our data further reveals a correlation between elevated CAMKV expression and adverse patient outcomes in NB. RNA sequencing and mass spectrometry analyses indicate that *CAMKV* knockdown leads to extensive transcriptomic and proteomic changes in NB cells. Moreover, we identified small molecules capable of inhibiting CAMKV activity, effectively suppressing tumor growth both in vitro and in a mouse NB xenograft model. These findings suggest that CAMKV serves as a potential biomarker for NB prognosis and a novel therapeutic target for NB treatment.

Elevated CREB expression and phosphorylation are commonly observed in various cancers. In our study, we discovered that CREB is highly phosphorylated at Ser133 in NB cells, contributing to NB cell proliferation. Additionally, upon CAMKV knockdown in NB cells, we noted the downregulation of many established CREB target genes (table S6). These genes are known to play crucial roles in neuronal system development and cancer progression (*56*). For instance, SCG2 exhibits high expression in the brain, adrenal gland, pituitary gland, and NB cells, regulating neuronal differentiation and safeguarding NB cells from nitric oxide-induced apoptosis (*77*). Similarly, BDNF and its receptor, TrkB, crucial for neuronal cell survival and development, are highly expressed in NB tumors, promoting resistance to genotoxic stress (*78–80*). This implies that CAMKV serves as a central hub controlling the expression of the genes that promote NB cell proliferation.

CREB has been reported to be a substrate for members of the CAMK2 subfamily (CAMK2A, CAMK2B, CAMK2D, and CAMK2G) and CAMK4 in many cell types (*28*, *29*). Intriguingly, the knockout of any of these CaMK2 members or CAMK4 did not notably inhibit CREB phosphorylation or NB cell proliferation (Fig. 1, D and E). Unexpectedly, we also observed that elevated expression of CaMK2A, CaMK2B, CAMK2D, CAMK2G, and CAMK4 in NB primary tumor specimens correlated with improved patient outcomes (Fig.S12). Further investigation is required to elucidate the role of other CaMK members in NB tumorigenesis.

A prior study claimed that recombinant CAMKV purified from bacteria lacks kinase activity, suggesting CAMKV is a non-functional pseudokinase (*30*). Another study proposed that CAMKV is vital for synaptic transmission and plasticity and maintaining dendritic spine structure, but no kinase activity is required for this function (*31*). We demonstrated for the first time that CAMKV kinase activity is required for NB tumor growth. It is plausible that CAMKV purified in a prokaryotic expression system lacks the correct folding or post-translational modifications necessary for its kinase activity. Furthermore, we identified threonine 183 (Thr183) as a critical site for CAMKV kinase activation and function in NB cells. However, further investigation is needed to uncover the upstream signaling pathway(s) that lead to CAMKV phosphorylation and activation in NB cells.

Calmodulin binding and subsequent autophosphorylation play crucial roles in regulating CaMKs’ activation (*81*). The interaction between calmodulin and the kinase is a vital regulatory step, promoting autophosphorylation and initiating the activation cascade. Once the critical amino acid is phosphorylated, it disrupts the autoinhibitory domain, which allows the kinase to establish a self-sustaining activation and render calmodulin dispensable for maintaining kinase activity (*81*, *82*). Consistent with these early reports, we found that the calmodulin binding domain is required for the initial activation of CAMKV but is not required for maintaining its kinase activity. To avoid artificial activations of CAMKV in vitro, we did not add calmodulin in our in vitro kinase assay to examine the kinase activity of endogenously activated CAMKV in SK-N-AS cells.

Alternative splicing contributes to transcript variation, proteome diversity, and the specificity of numerous cellular processes (*83–85*). Aberrant splicing events are also common in NB cells and contribute to cancer progression (*86–89*). Notably, four CaMKII genes (CAMK2A, CAMK2B, CAMK2G, and CAMK2D) undergo alternative splicing, which results in changes in specific kinase activity and differential responses to calcium stimulation in vitro (*90*, *91*). In this study, two isoforms of CAMKV with kinase activity were identified in NB cells. The short isoform exhibited higher kinase activity than the full-length isoform, suggesting that the deleted region in the short isoform negatively regulates CAMKV kinase activity. Further investigation is needed to determine whether the alternative splicing of *CAMKV* contributes to NB development.

*MYCN* amplification is a powerful tool for stratifying NB patients and predicting their prognosis (*3*, *4*). However, it occurs in less than half of high-risk NB patients, indicating that more than half still experience poor outcomes despite the absence of *MYCN* amplification (*7*, *8*). This underscores the need for additional markers to enhance risk stratification. We found that NB patients with high *CAMKV* expression have significantly worse survival than those with lower expression, suggesting that elevated *CAMKV* levels are at least partially correlated with a more aggressive NB phenotype. Consistent with a recent report (*34*), our study also demonstrated that *CAMKV* is transcriptionally regulated by MYCN/MYC. Therefore, patients with high-risk NB without *MYCN* amplification could potentially be identified by this novel marker, CAMKV, supplementing the limitations of *MYCN* as a sole stratification tool. However, additional studies are needed to further investigate the association between CAMKV and MYCN/MYC protein expression levels in clinical specimens.

Our RNA-seq and mass spectrometry data revealed extensive transcriptomic, proteomic, and phosphorylation changes in NB cells upon *CAMKV* knockdown. These changes included phosphorylation of eight transcription factors. Among them, we identified two decreased phosphopeptides from the transcription factor GATA2 at Ser182 and Ser192, implying that GATA2 is another CAMKV substrate. Like *CAMKV*, *GATA2* exhibits specific overexpression in NB cells and may play a critical role in mediating CAMKV kinase signaling. Using an anti-phospho-GATA2 (Ser192) antibody, we confirmed its decreased phosphorylation in CAMKV-knockdown NGP cells while the total GATA2 protein level remained unchanged (Fig.S10A). Besides transcription factors, we also observed decreased phosphorylation of several kinases after CAMKV knockdown (table S8 and S9). One example is STK10 (Serine/Threonine-Protein Kinase 10), expressed in various tumor cell lines and highly proliferative tissues (*92*). STK10 plays a crucial role in tumor progression and its knockdown promotes tumor cell apoptosis (*93*, *94*). Another example is RIOK1 (RIO kinase 1), which is known to promote cancer cell proliferation and invasion (*95*). These findings suggest that CAMKV regulates the phosphorylation of multiple categories of proteins to promote NB growth.

K252a has been shown to exert inhibitory effects on various signaling pathways by suppressing the activity of multiple kinases including TrkA and TrkB tyrosine kinases (*96–101*). Therefore, K252a likely inhibits other kinase-mediated pathways contributing to NB cell proliferation and tumor growth besides CAMKV. However, all the candidate inhibitors we identified are not specific for CAMKV and screening specific CAMKV kinase inhibitors is therefore needed in the future. Our current results provide proof-of-concept that inhibiting CAMKV activity with small molecule inhibitors represents a promising strategy for developing novel NB therapies.

In this study, we demonstrated that MYCN/MYC transcriptionally up-regulates *CAMKV* expression. Once activated, CAMKV directly phosphorylates CREB and other targets to modulate gene expression, thereby enhancing NB cell proliferation. Inactivating the *CAMKV* gene or inhibiting its kinase activity with small molecule inhibitors suppresses tumor growth, which suggests that CAMKV kinase represents a novel therapeutic avenue and is a potential prognostic marker for NB.

## Author contributions

Conceptualization: JY. Methodology: LLW, JW, SYJ, CZ, JY. Investigation: YY, YZ, ZS, FC, LLW, JMC, KL, DS, DQ, JW, SA, CZ. Visualization: YY, YZ, ZS, FC, LLW, JMC, KL, JW. Funding acquisition: JY. Project administration: JY. Supervision: CZ, JY. Writing – original draft: YY, YZ, ZS, FC, LLW, SA, CZ, JY. Writing – review & editing: JW, SA, BRD, JSD, MF, JSY, EW, JY.

## Data and materials availability

All data associated with this study are present in the paper or the Supplementary Materials. RNA-seq data are available at GEO under accession number GSE255691(CAMKV knockdown) and GSE255692 (OTSSP167 treatment). Proteomics data have been deposited to the ProteomeXchange Consortium via the PRIDE with accession number PXD049345 (global proteome) and PXD049413 (phosphoproteomics).

---

**Supplementary figure 1.**
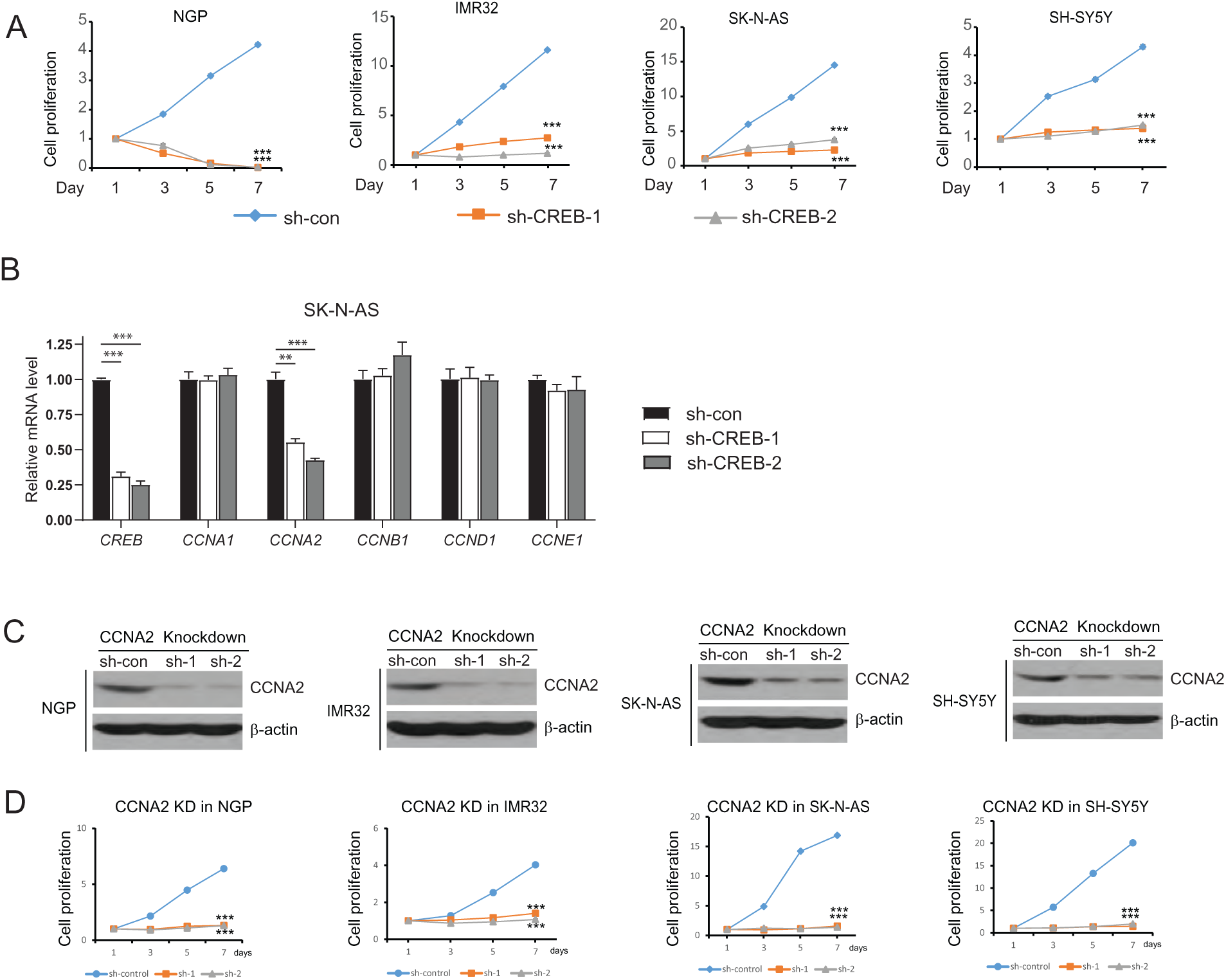
The effects of *CREB* and *CCNA2* gene knockdown on cell proliferation in NB cells. **(A)** The effects of *CREB* knockdown on NB cell proliferation. NB cells with *CREB* knockdown or control shRNA were seeded in 96-well plates at a concentration of 1 × 10^4^ cells per well. CCK8 assay was performed on day three to determine the cell viability. Experiment was performed in triplicate. **(B)** The effects of *CREB* knockdown on the expression of different cyclins in SK-N-AS cell. The mRNA levels were detected by real-time PCR. **(C, D)** The effects of *CCNA2* knockdown on NB cell proliferation. NB cells were transduced with shRNAs targeting *CCNA2* or with control shRNA. The protein levels of CCNA2 and β-actin in the cell lysates were detected by immunoblotting.

**Supplementary figure 2.**
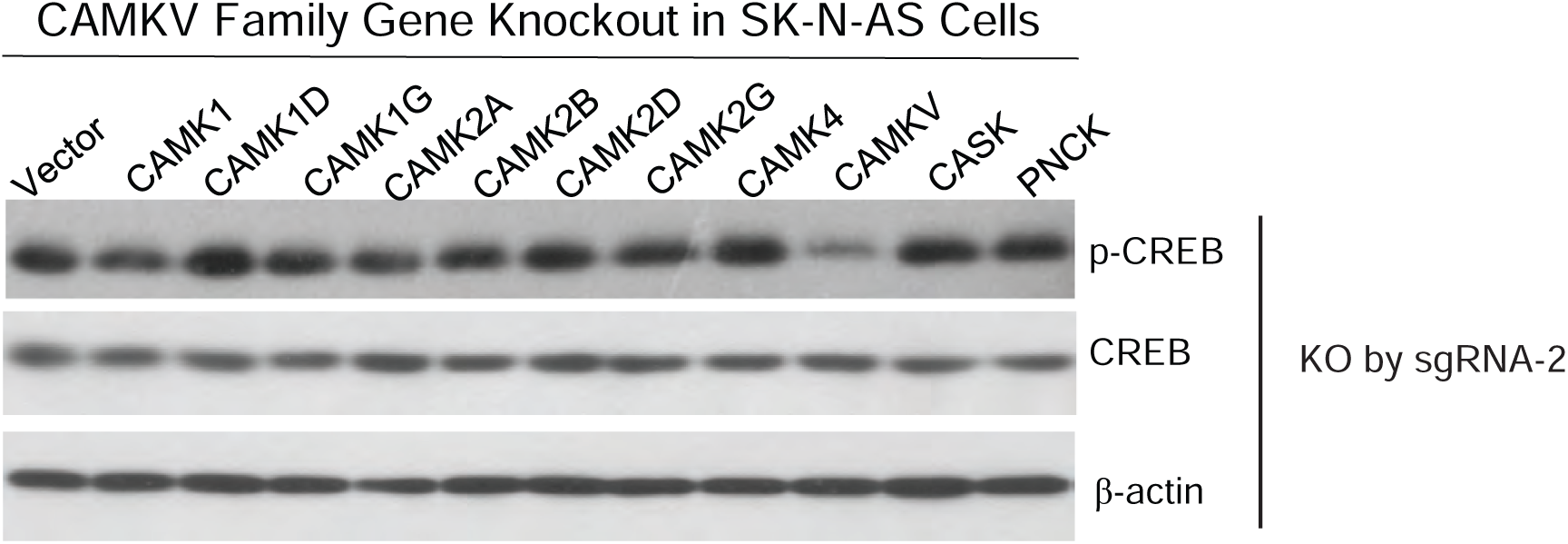
The effects of CRISPR/Cas9 knockout of each member of Ca2+/calmodulin-dependent kinases (CaMKs) on CREB phosphorylation by the second set of sgRNAs.

**Supplementary figure 3.**
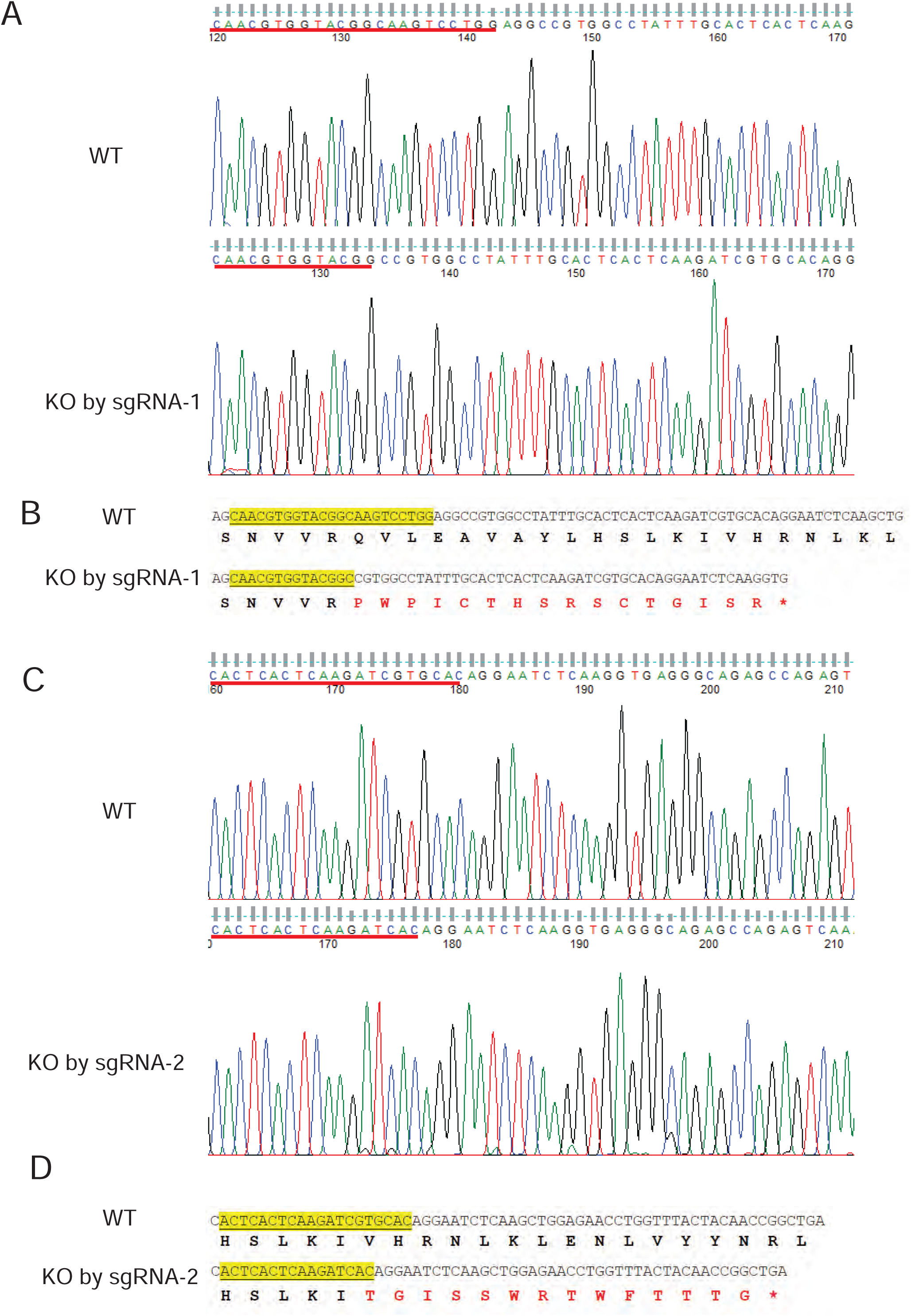
Sequencing results of CAMKV knockout in SK-N-AS cells. **(A, C)** Sequencing results of single clones from CAMKV knockout mediated by guide RNA-1 (A) and guide RNA-2 (C). **(B, D)** The reading frame-shifted CAMKV genomic sequences after genome editing by guide RNA-1 (B) and guide RNA-2 (D).

**Supplementary figure 4.**
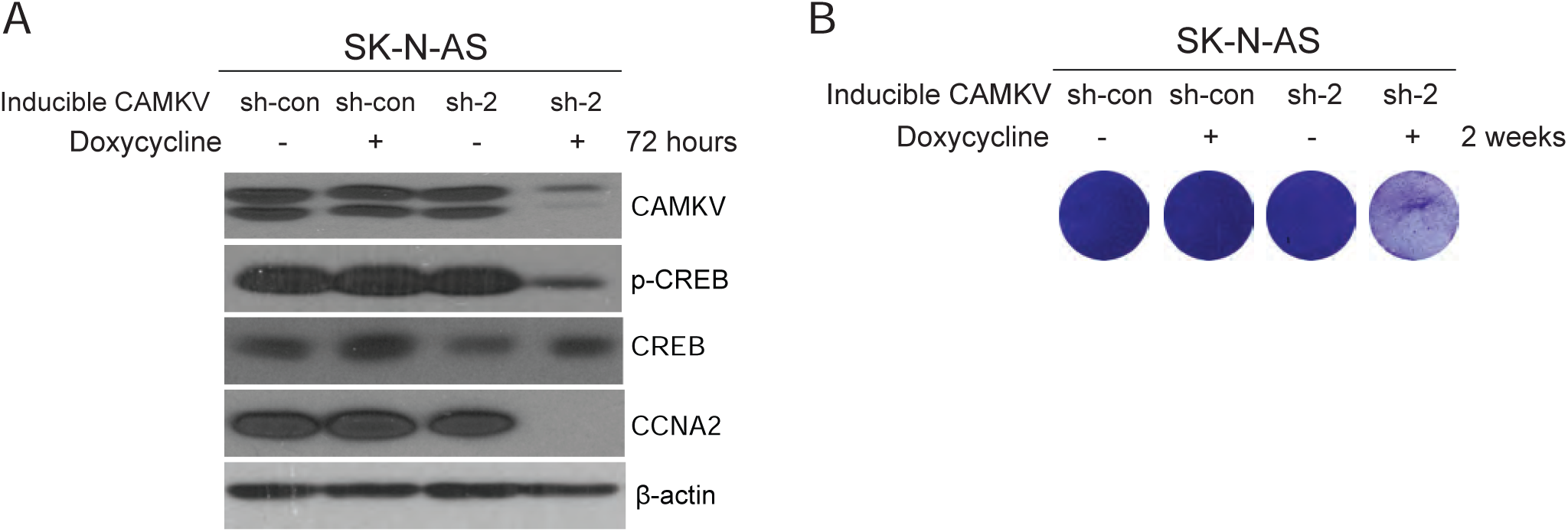
Effects of CAMKV inducible knockdown mediated by an inducible piggyBac (PB) system in SK-N-AS cells. **(A)** The effects of inducible knockdown of *CAMKV* gene on CREB phosphorylation. SK-N-AS cells were treated with doxycycline (100 ng/mL, 72 hours) and harvested for immunoblotting analysis. **(B)** SK-N-AS cells were treated with doxycycline (100 ng/mL, 14 days) and stained with crystal violet.

**Supplementary figure 5.**
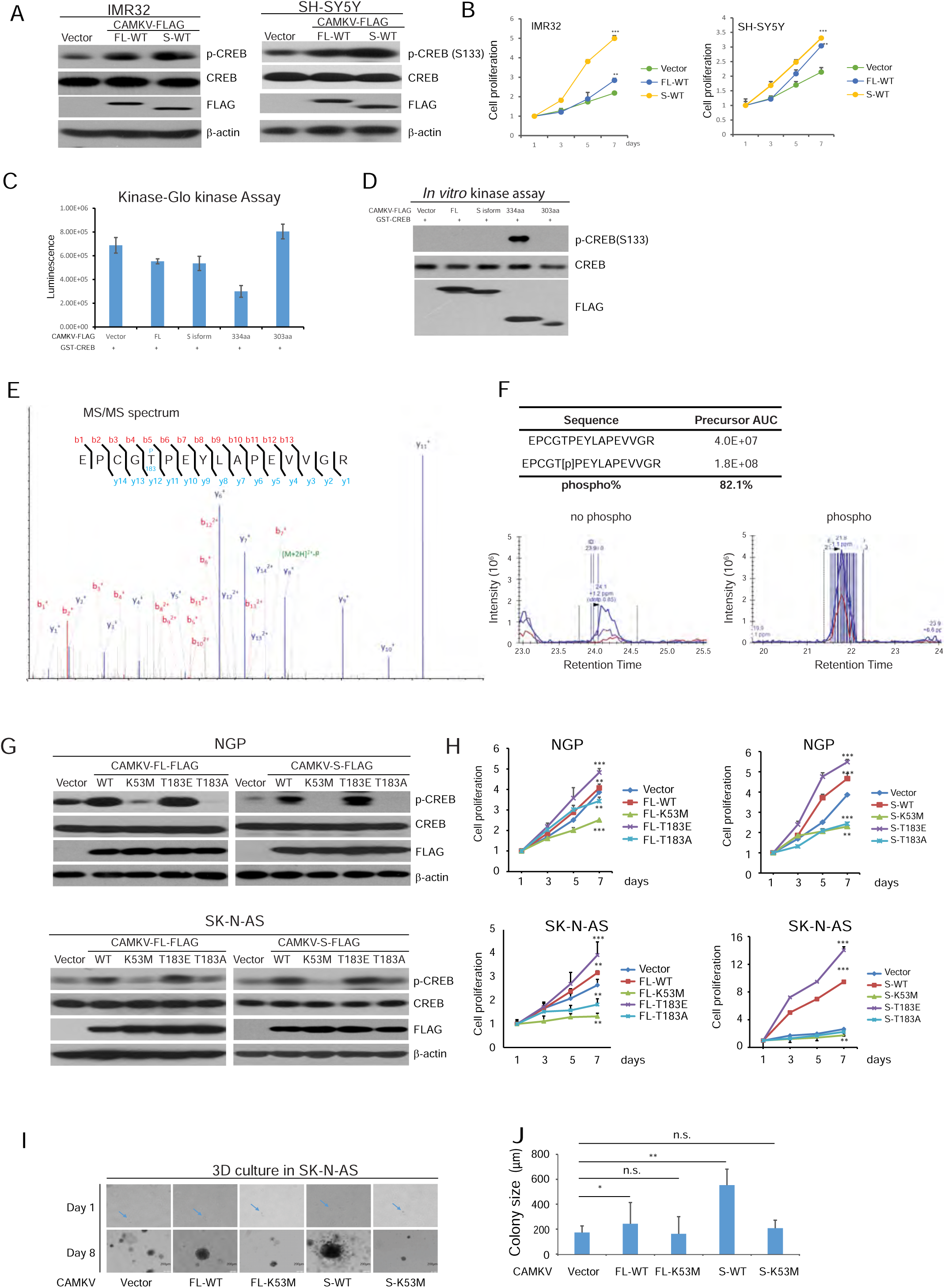
The kinase activity of CAMKV is required for NB cell proliferation and phosphorylation at Thr183 is crucial for kinase activation. **(A, B)** The effects of over-expression of CAMKV-FL-WT or CAMKV-S-WT on CREB phosphorylation (A) and cell proliferation (B) of IMR32 and SH-SY5Y cells. **(C, D)** In vitro phosphorylation of recombinant GST-CREB by CAMKV-FL, CAMKV-S, CAMKV-334aa, and CAMKV-303aa purified from overexpressed HEK293T cells by Kinase-Glo® luminescent kinase assay (C) and in vitro kinase assay (D). **(E, F)** Identification of CAMKV phosphorylation site by mass spectrometry (MS). (E) The representative MS/MS spectrum of the tryptic phospho-peptide EPCGT(p)PEYLAPEVVGR (p denotes phosphorylation), which identified phosphorylation at Thr-183, is shown. (F) The percent of the phospho-peptide from MS analysis. **(G, H)** Overexpressing either the FL mutants (left) or S mutants (right), the threonine 183 residue is the critical phosphorylation site for optimal CAMKV function to promote CREB phosphorylation (G) and NB cell proliferation (H) in NGP and SK-N-AS cell lines. **(I, J)** The effects of overexpression of CAMKV WT or mutants on colony formation of SK-N-AS cells in Matrigel. The blue arrows point single cells, and the scale indicates 200 µm. (I) Colony images were captured at 100 × magnification under a microscope. (J) The sizes of twenty colonies from each well were measured, averaged, and presented as mean ± S.D.

**Supplementary figure 6.**
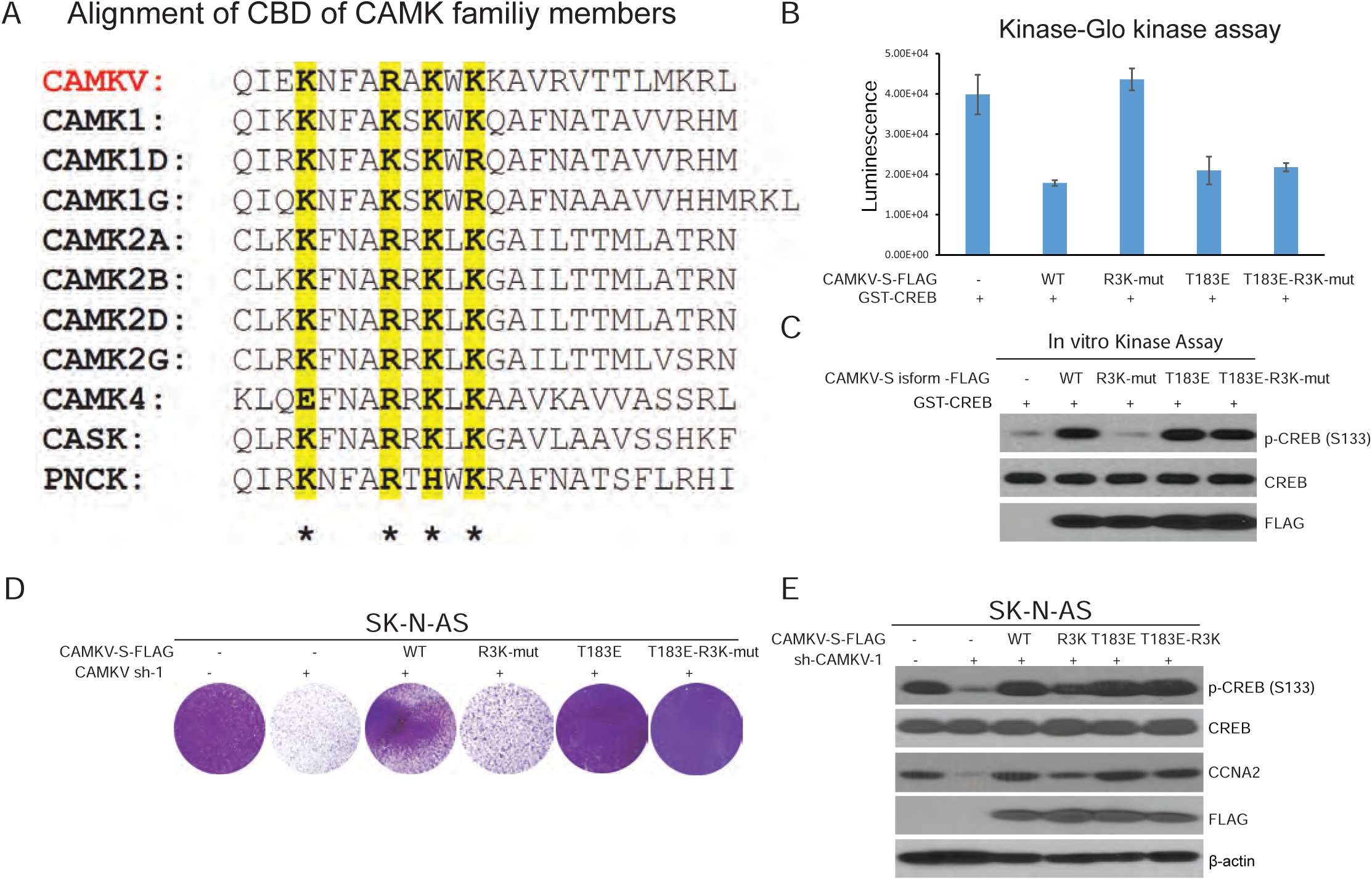
Calmodulin binding domain is required for the initial activation of CAMKV kinase in NB cells. **(A)** Sequence alignment of calmodulin binding domain (CBD) of CAMK family members with conserved lysines (K) and arginines (R) highlighted in yellow. **(B, C)** In vitro phosphorylation of recombinant GST-CREB by CAMKV-S-WT, CAMKV-S-R3K-mut, CAMKV-S-T183E and CAMKV-S-T183E-R3K-mut. by Kinase-Glo® luminescent kinase assay (B) and an in vitro kinase assay (C). **(D, E)** The effects of stable expression of shRNA resistant CAMKV-S-WT, -S or their mutants on cell proliferation (D) and CREB phosphorylation (E) in SK-N-AS cells, in which endogenous CAMKV was knocked down.

**Supplementary figure 7.**
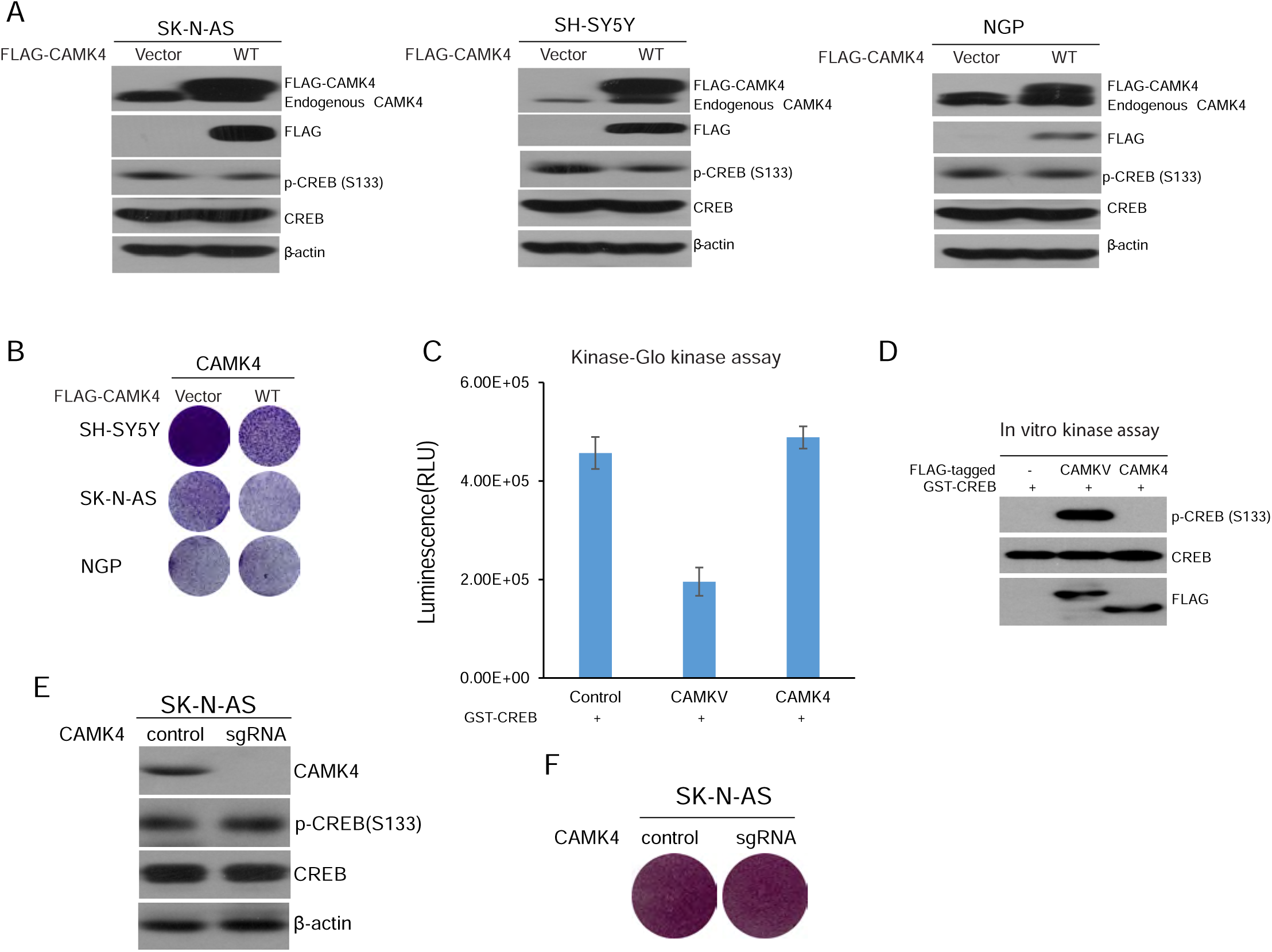
CAMK4 is not required for CREB phosphorylation in NB cells and NB cell proliferation. **(A)** The effects of CAMK4 overexpression on total and phospho-CREB by immunoblotting in SK-N-AS, SH-SY5Y, and NGP cell lines. beta-actin was used as a loading control. by immunoblotting. **(B)** The effects of CAMK4 overexpression on the NB cell proliferation by crystal violet staining after 7 days of culture. **(C, D)** In vitro phosphorylation of recombinant GST-CREB by FLAG-tagged CAMKV-FL compared to CAMK4-FL by a Kinase-Glo® luminescent kinase assay (C) and an in vitro kinase assay (D). **(E, F)** The effects of CAMK4 knockout on CREB phosphorylation (E) and cell proliferation (F) in SK-N-AS cells.

**Supplementary figure 8.**
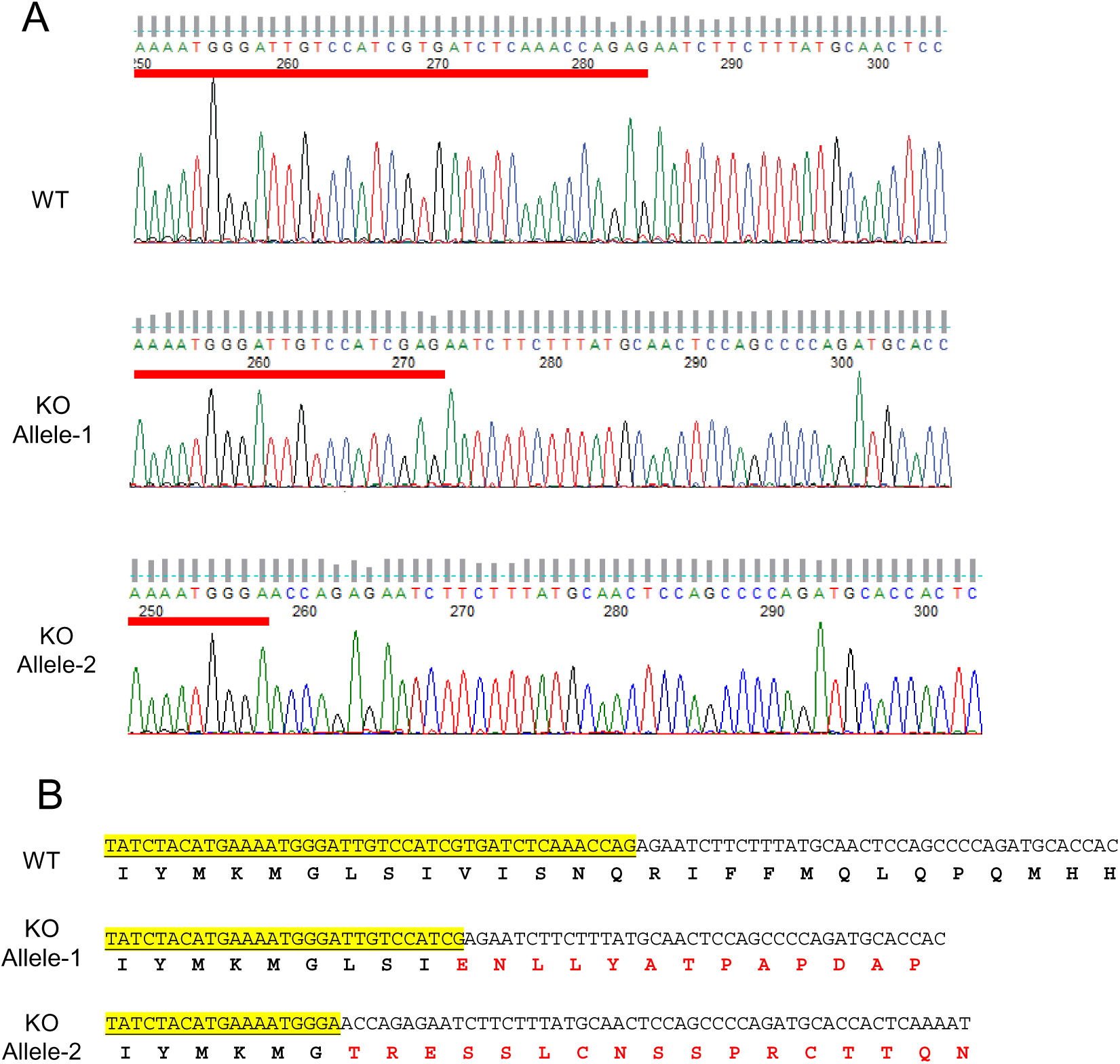
Sequencing results of CAMK4 knockout in SK-N-AS cells. **(A)** Sequencing results of single clones from CAMK4 knockout mediated by guide RNA-1. **(B)** The reading frame-shifted CAMK4 genomic sequences after genome editing by guide RNA-1.

**Supplementary figure 9.**
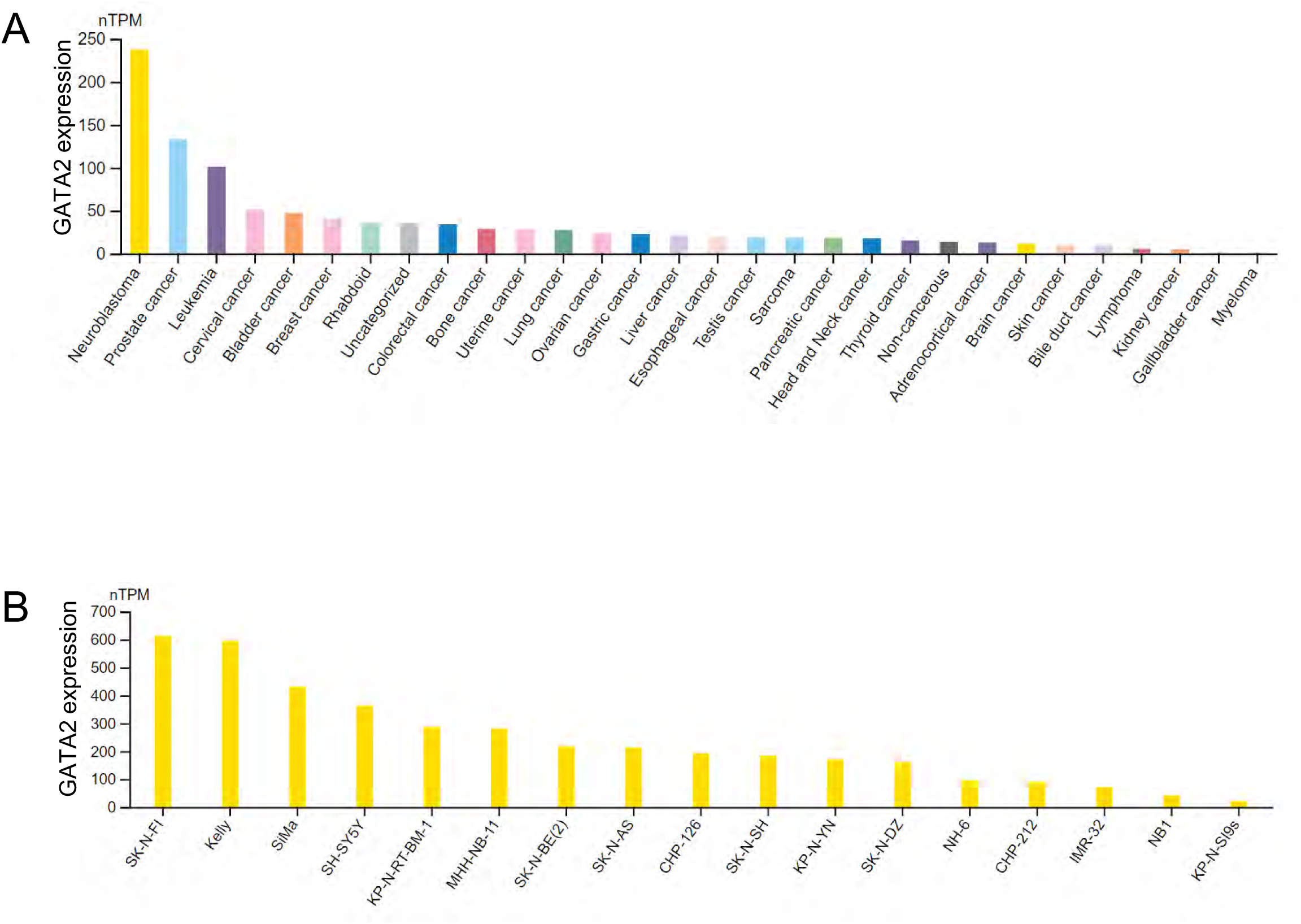
GATA2 is highly expressed in NB cells. **(A)** GATA2 mRNA level in different types of cancer cell lines. **(B)** GATA2 mRNA level in different NB cell lines. (https://www.proteinatlas.org/ENSG00000179348-GATA2/cell+line#neuroblastoma)

**Supplementary figure 10.**
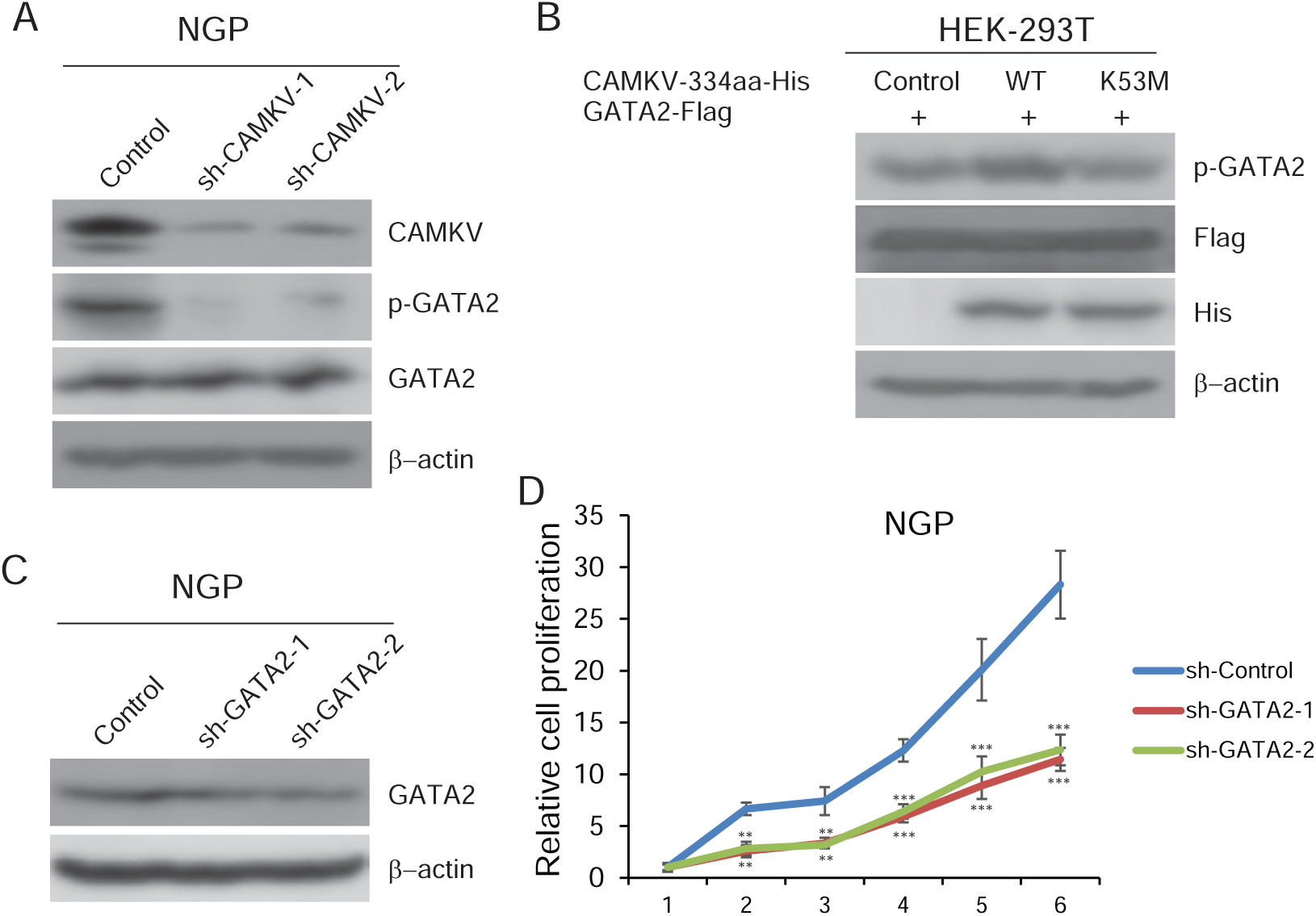
The phosphorylation of GATA2 is regulated by CAMKV. **(A)** The effect of CAMKV knockdown on the phosphorylation level of GATA2 in NGP cells. **(B)** The effect of overexpression of His-tagged CAMKV-334aa WT and K53M mutant on the phosphorylation level of co-transfected FLAG-GATA2 in HEK293T cells. **(C)** GATA2 protein level in GATA2-knockdown NGP cells. **(D)** The effect of GATA2 knockdown on NGP cell proliferation.

**Supplementary figure 11.**
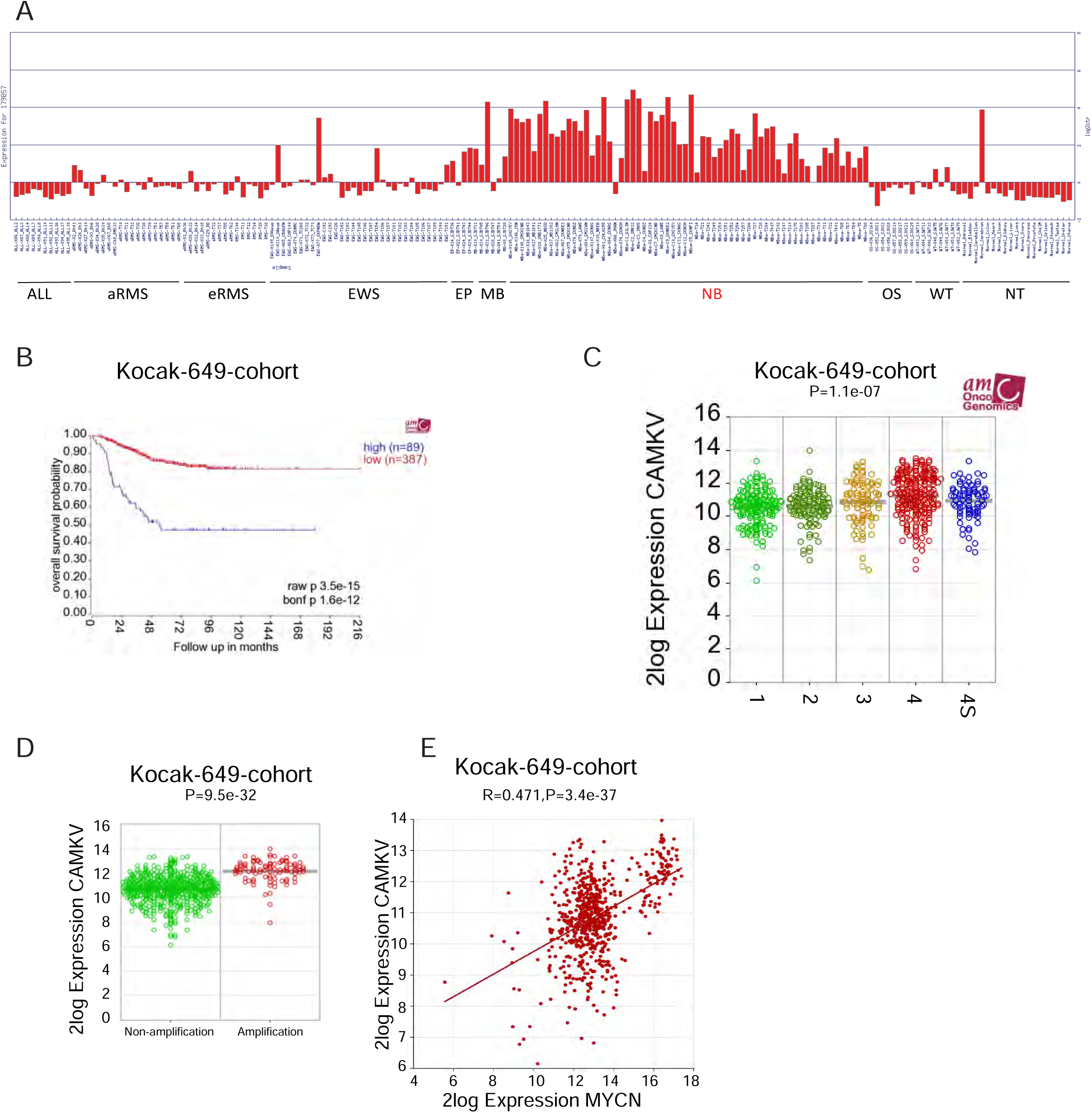
Correlation of high expression of *CAMKV* in tumors with a poor NB patient survival. **(A)** Relative *CAMKV* expression levels in a normal tissue and pediatric malignancy microarray array database, including acute lymphocytic leukemia (ALL), alveolar (aRMS) and embryonal (eRMS) rhabdomyosarcoma, Ewing’s sarcoma (EWS), ependymoma (EP), medulloblastoma (MB), neuroblastoma (NB), osteosarcoma (OS), Wilms’ tumor (WT), and normal tissue (NT). **(B)** Correlation of *CAMKV* mRNA levels and the overall survival of NB patients was analyzed in Kocak-649-cohort dataset. **(C, D)** *CAMKV* mRNA levels among different stages (Stage 1, n=153; Stage 2, n=113; Stage 3, n=91; Stage 4, n=214; Stage 4s, n=78) (C) and in samples with *MYCN* non-amplification (n=550) or amplification (n=93) (D). **(E)** Correlation of *CAMKV* and *MYCN* mRNA levels in all 649 NB tumor tissues.

**Supplementary figure 12.**
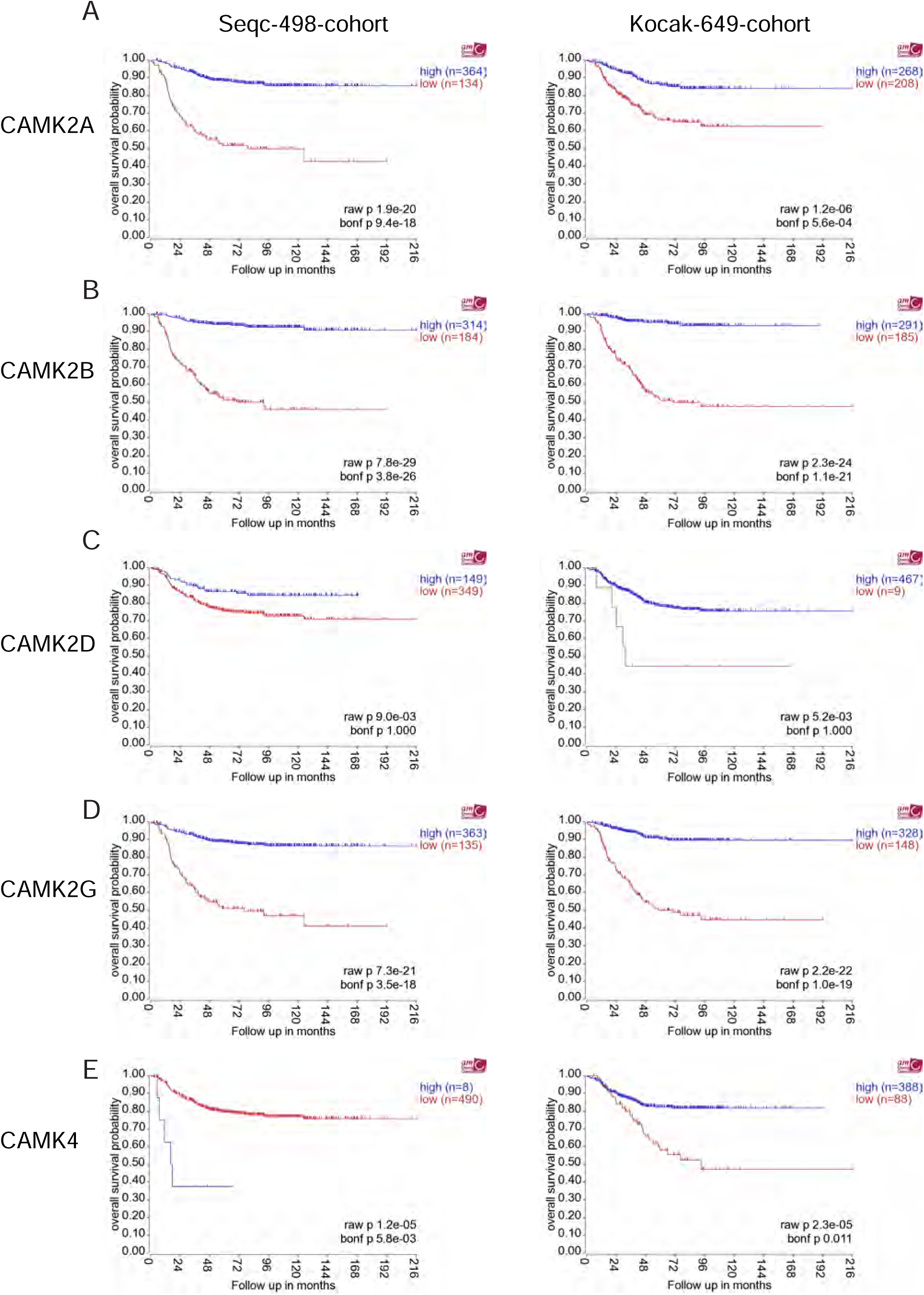
Correlations of the CAMK family members mRNA levels and patient overall survival. Correlations of *CAMK2A* (A), *CAMK2B* (B), *CAMK2D* (C), *CAMK2G* (D), or *CAMK4* (E) mRNA levels and patient overall survival in Seqc-498-cohort (left) and Kocak-649-cohort (right) datasets.

**Supplementary figure 13.**
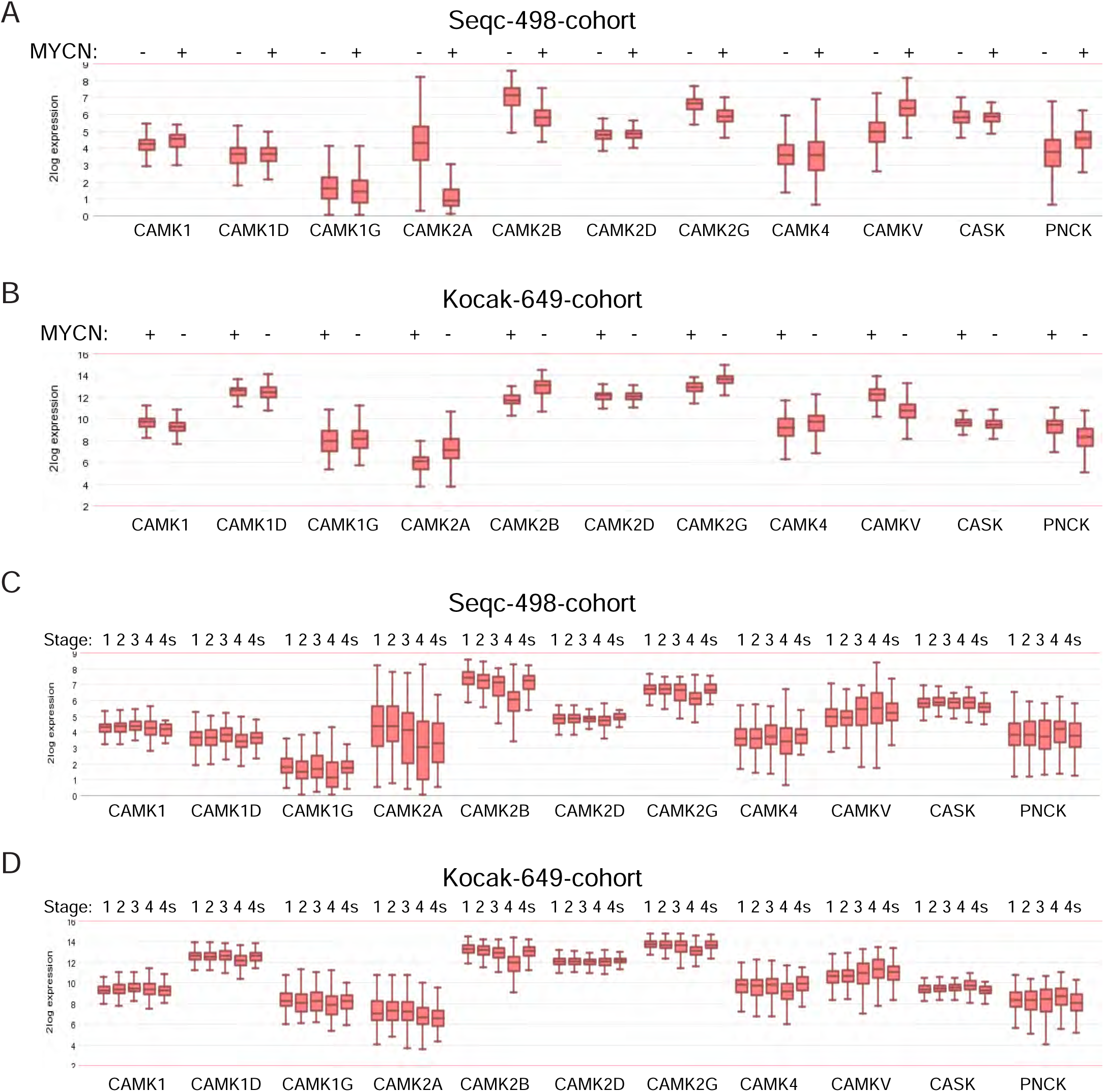
The mRNA levels of CAMK family members in different *MYCN* status and INSS stages. **(A, B)** The mRNA levels of *CaMKs* in NB tumor tissues between *MYCN* non-amplified and amplified groups in Seqc-498-cohort (A) and the Kocak-649-cohort datasets (B). **(C, D)** The mRNA levels of *CaMKs* in NB tumor tissues among different INSS stages from Seqc-498-cohort (C) and the Kocak-649-cohort (D) datasets.

**Supplementary figure 14.**
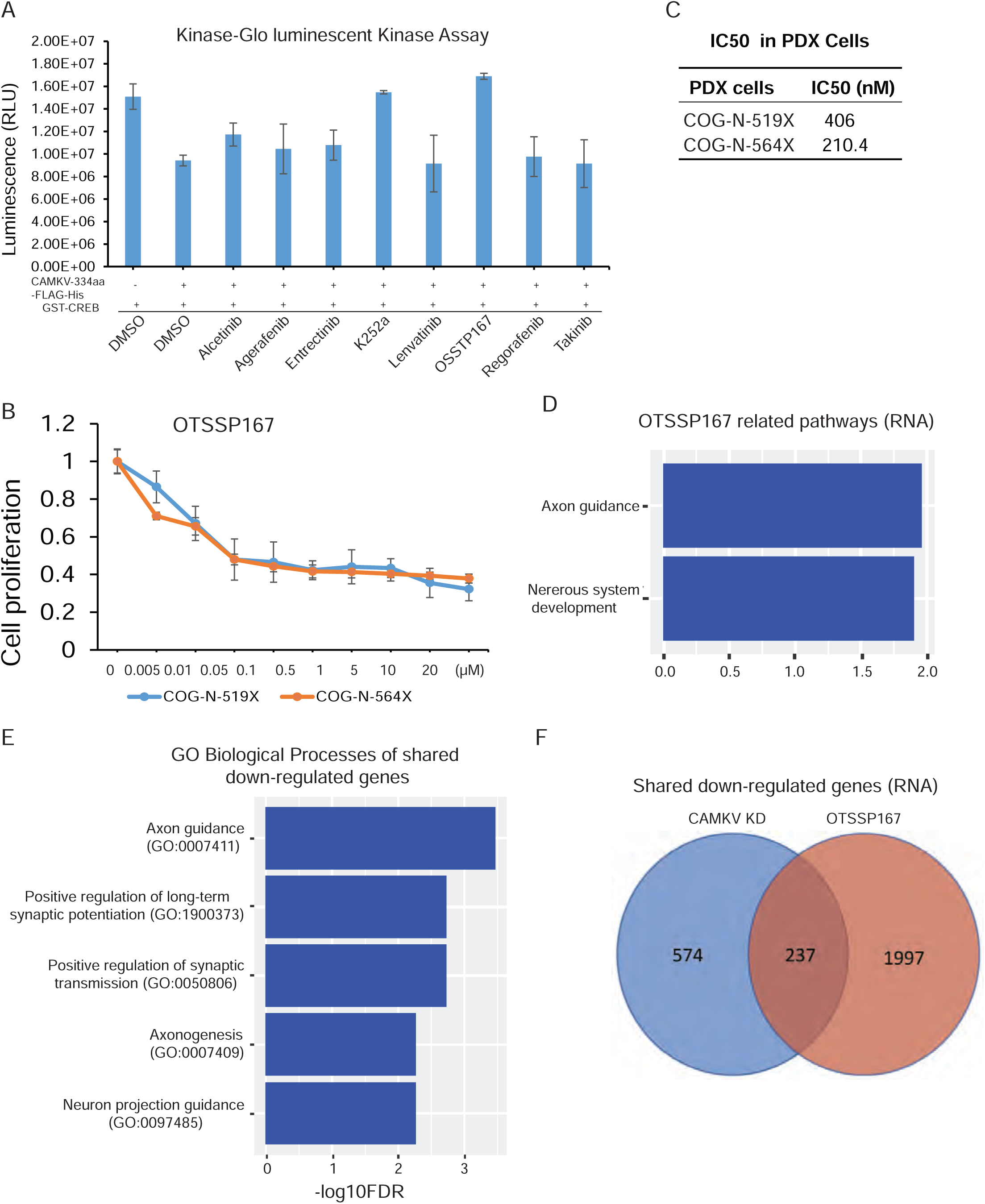
OTSSP167 inhibits CAMKV kinase activity in NB cells. **(A)** The effects of Alcetinib, Agerafenib, Entrectinib, K252a, Lenvatinib, OTSSP167, Regorafenib, and Takinib on ATP consumption of CAMKV-334a incubated with GST-CREB in the Kinase-Glo Assay. **(B)** The effect of OSTSSP167 ex vivo treatment on cell proliferation of NB PDX cells. **(C)** The IC_50_ value of OTSSP167 in NB PDX cells, COG-N-519X and COG-N-564X. **(D)** Gene set enrichment analysis of downregulated genes in OTSSP167 treated NGP cells showing enrichment of neuronal-related pathways. **(E, F)** Integrated analysis of RNA-seq results between CAMKV knockdown and OTSSP167 treated NGP cells.

**Supplementary figure 15.**
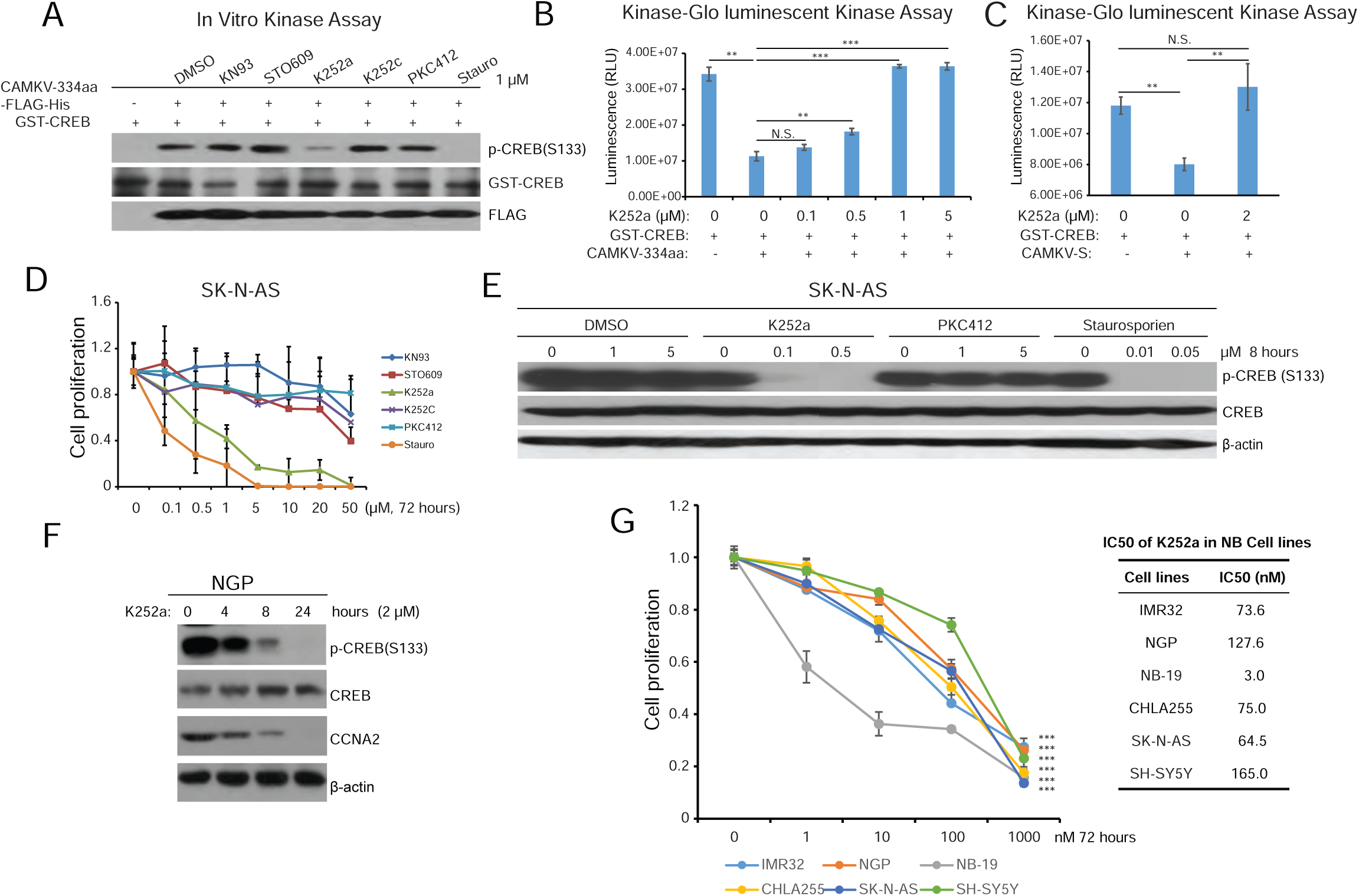
K252a inhibits CAMKV kinase-mediated CREB phosphorylation. **(A)** The effects of CAMK-family inhibitors KN93, STO609, K252a, K252c, PKC412, and staurosporine (Stauro) on CAMKV-334aa-mediated phosphorylation of recombinant GST-CREB in an in vitro kinase assay. **(B, C)** K252a inhibited ATP consumption for recombinant GST-CREB phosphorylation by the purified CAMKV-334aa (B) or CAMKV-S (C) in the Kinase-Glo Assay. **(D)** The effects of CAMK family member inhibitors on SK-N-AS cell viability. **(E)** The effects of K252a, PKC412, and staurosporine (Stauro) on CREB phosphorylation level in NB cells. **(F)** The effects of K252a on CREB phosphorylation and CCNA2 expression in NB cells. **(G)** Inhibitory effects of K252a on the viability of different NB cell lines.

**Supplementary figure 16.**
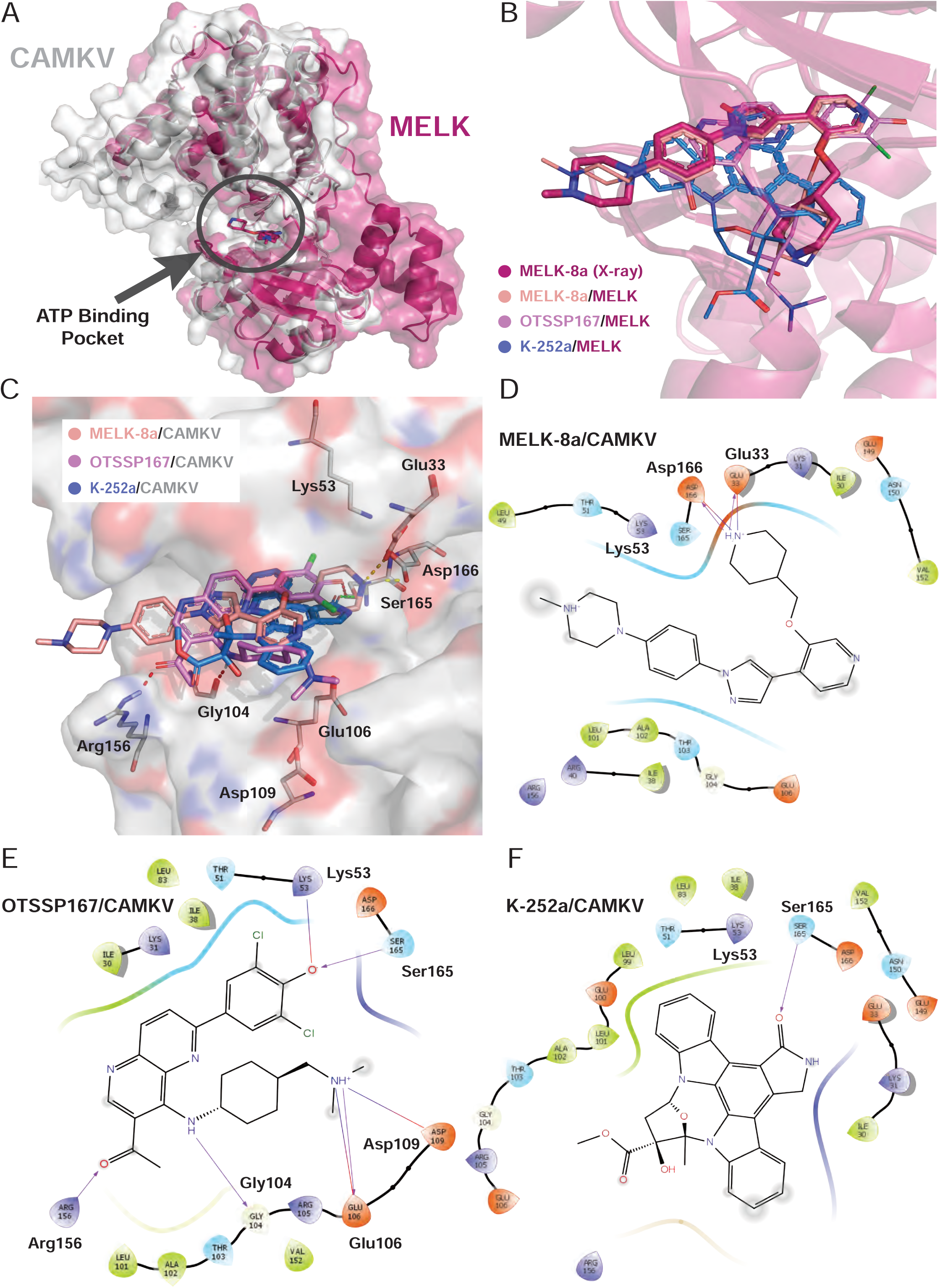
Molecular docking of MELK-8a, OTSSP167, and K-252a with CAMKV and MELK. **(A)** Alignment between CAMKV (white, generated by AlpahFold2) and MELK-8a/MELK (warm pink, PDB ID: 5IH9), and the ATP binding pocket was highlighted. **(B)** The superposition of MELK-8a redock pose (wheat) and its original inhibitor (warm pink), and the docking poses of OTSSP167 (violet) and K-252a (marine) within the ATP-binding pocket of MELK. **(C)** The docking poses of MELK-8a, OTSSP167, and K-252a within the ATP-binding pocket of CAMKV. **(D-F)** The 2D-interactive patterns between key residues and MELK-8a (D), OTSSP167 (E), and K-252a (F) within the ATP-binding pocket of CAMKV. Molecular docking was performed by Glide with XP precision; 2D interactive pattern was illustrated by Ligand Interactive Diagram; 3D binding mode was illustrated by PyMOL.

## References

1. J. R. Park, R. Bagatell, W. B. London, J. M. Maris, S. L. Cohn, K. K. Mattay, M. Hogarty, COG Neuroblastoma Committee, Children’s Oncology Group’s 2013 blueprint for research: neuroblastoma. Pediatr Blood Cancer 60, 985–993 (2013).

2. G. M. Brodeur, Neuroblastoma: biological insights into a clinical enigma. Nat Rev Cancer 3, 203–216 (2003).

3. G. M. Brodeur, R. C. Seeger, M. Schwab, H. E. Varmus, J. M. Bishop, Amplification of N-myc in untreated human neuroblastomas correlates with advanced disease stage. Science 224, 1121–1124 (1984).

4. M. Brockmann, E. Poon, T. Berry, A. Carstensen, H. E. Deubzer, L. Rycak, Y. Jamin, K. Thway, S. P. Robinson, F. Roels, O. Witt, M. Fischer, L. Chesler, M. Eilers, Small molecule inhibitors of aurora-a induce proteasomal degradation of N-myc in childhood neuroblastoma. Cancer Cell 24, 75–89 (2013).

5. A. Pession, R. Tonelli, R. Fronza, E. Sciamanna, R. Corradini, S. Sforza, T. Tedeschi, R. Marchelli, L. Montanaro, C. Camerin, M. Franzoni, G. Paolucci, Targeted inhibition of NMYC by peptide nucleic acid in N-myc amplified human neuroblastoma cells: cell-cycle inhibition with induction of neuronal cell differentiation and apoptosis. Int J Oncol 24, 265–272 (2004).

6. A. Puissant, S. M. Frumm, G. Alexe, C. F. Bassil, J. Qi, Y. H. Chanthery, E. A. Nekritz, R. Zeid, W. C. Gustafson, P. Greninger, M. J. Garnett, U. McDermott, C. H. Benes, A. L. Kung, W. A. Weiss, J. E. Bradner, K. Stegmaier, Targeting MYCN in neuroblastoma by BET bromodomain inhibition. Cancer Discov 3, 308–323 (2013).

7. X. Liu, P. Mazanek, V. Dam, Q. Wang, H. Zhao, R. Guo, J. Jagannathan, A. Cnaan, J. M. Maris, M. D. Hogarty, Deregulated Wnt/beta-catenin program in high-risk neuroblastomas without MYCN amplification. Oncogene 27, 1478–1488 (2008).

8. F. Westermann, D. Muth, A. Benner, T. Bauer, K.-O. Henrich, A. Oberthuer, B. Brors, T. Beissbarth, J. Vandesompele, F. Pattyn, B. Hero, R. König, M. Fischer, M. Schwab, Distinct transcriptional MYCN/c-MYC activities are associated with spontaneous regression or malignant progression in neuroblastomas. Genome Biol 9, R150 (2008).

9. M. W. Zimmerman, Y. Liu, S. He, A. D. Durbin, B. J. Abraham, J. Easton, Y. Shao, B. Xu, S. Zhu, X. Zhang, Z. Li, N. Weichert-Leahey, R. A. Young, J. Zhang, A. T. Look, MYC Drives a Subset of High-Risk Pediatric Neuroblastomas and Is Activated through Mechanisms Including Enhancer Hijacking and Focal Enhancer Amplification. Cancer Discov 8, 320–335 (2018).

10. M. J. Henley, A. N. Koehler, Advances in targeting “undruggable” transcription factors with small molecules. Nat Rev Drug Discov 20, 669–688 (2021).

11. P. K. Brindle, M. R. Montminy, The CREB family of transcription activators. Curr Opin Genet Dev 2, 199–204 (1992).

12. H. Bito, K. Deisseroth, R. W. Tsien, CREB phosphorylation and dephosphorylation: a Ca(2+)-and stimulus duration-dependent switch for hippocampal gene expression. Cell 87, 1203–1214 (1996).

13. L. W. Yuan, J. E. Gambee, Histone acetylation by p300 is involved in CREB-mediated transcription on chromatin. Biochim Biophys Acta 1541, 161–169 (2001).

14. R. A. Woo, R. Y. C. Poon, Cyclin-dependent kinases and S phase control in mammalian cells. Cell Cycle 2, 316–324 (2003).

15. S. Dalle, J. Quoyer, E. Varin, S. Costes, Roles and regulation of the transcription factor CREB in pancreatic β -cells. Curr Mol Pharmacol 4, 187–195 (2011).

16. S. Boulon, J.-C. Dantonel, V. Binet, A. Vié, J.-M. Blanchard, R. A. Hipskind, A. Philips, Oct-1 potentiates CREB-driven cyclin D1 promoter activation via a phospho-CREB- and CREB binding protein-independent mechanism. Mol Cell Biol 22, 7769–7779 (2002).

17. T. Mabuchi, K. Kitagawa, K. Kuwabara, K. Takasawa, T. Ohtsuki, Z. Xia, D. Storm, T. Yanagihara, M. Hori, M. Matsumoto, Phosphorylation of cAMP response element-binding protein in hippocampal neurons as a protective response after exposure to glutamate in vitro and ischemia in vivo. J Neurosci 21, 9204–9213 (2001).

18. A. Riccio, S. Ahn, C. M. Davenport, J. A. Blendy, D. D. Ginty, Mediation by a CREB family transcription factor of NGF-dependent survival of sympathetic neurons. Science 286, 2358–2361 (1999).

19. M. R. Walton, I. Dragunow, Is CREB a key to neuronal survival? Trends Neurosci 23, 48– 53 (2000).

20. E. Ciani, S. Guidi, G. Della Valle, G. Perini, R. Bartesaghi, A. Contestabile, Nitric oxide protects neuroblastoma cells from apoptosis induced by serum deprivation through cAMP-response element-binding protein (CREB) activation. J Biol Chem 277, 49896–49902 (2002).

21. E.-M. Park, S. Cho, Enhanced ERK dependent CREB activation reduces apoptosis in staurosporine-treated human neuroblastoma SK-N-BE(2)C cells. Neurosci Lett 402, 190– 194 (2006).

22. N. Zhang, Q. Wen, L. Ren, W. Liang, Y. Xia, X. Zhang, D. Zhao, D. Sun, Y. Hu, H. Hao, Y. Yan, G. Zhang, J. Yang, T. Kang, Neuroprotective effect of arctigenin via upregulation of P-CREB in mouse primary neurons and human SH-SY5Y neuroblastoma cells. Int J Mol Sci 14, 18657–18669 (2013).

23. Y. M. Yang, L. R. Dolan, Z. Ronai, Expression of dominant negative CREB reduces resistance to radiation of human melanoma cells. Oncogene 12, 2223–2233 (1996).

24. P. I. Hanson, H. Schulman, Neuronal Ca2+/calmodulin-dependent protein kinases. Annu Rev Biochem 61, 559–601 (1992).

25. A. R. Means, Regulatory cascades involving calmodulin-dependent protein kinases. Mol Endocrinol 14, 4–13 (2000).

26. K. U. Bayer, H. Schulman, CaM kinase: Still inspiring at 40. Neuron 103, 380–394 (2019).

27. O. Rodriguez-Mora, M. M. LaHair, C. J. Howe, J. A. McCubrey, R. A. Franklin, Calcium/calmodulin-dependent protein kinases as potential targets in cancer therapy. Expert Opin Ther Targets 9, 791–808 (2005).

28. P. Sun, H. Enslen, P. S. Myung, R. A. Maurer, Differential activation of CREB by Ca2+/calmodulin-dependent protein kinases type II and type IV involves phosphorylation of a site that negatively regulates activity. Genes Dev 8, 2527–2539 (1994).

29. L. Racioppi, A. R. Means, Calcium/calmodulin-dependent kinase IV in immune and inflammatory responses: novel routes for an ancient traveller. Trends Immunol 29, 600–607 (2008).

30. M. Godbout, M. G. Erlander, K. W. Hasel, P. E. Danielson, K. K. Wong, E. L. Battenberg, P. E. Foye, F. E. Bloom, J. G. Sutcliffe, 1G5: a calmodulin-binding, vesicle-associated, protein kinase-like protein enriched in forebrain neurites. J Neurosci 14, 1–13 (1994).

31. Z. Liang, Y. Zhan, Y. Shen, C. C. L. Wong, J. R. Yates, F. Plattner, K.-O. Lai, N. Y. Ip, The pseudokinase CaMKv is required for the activity-dependent maintenance of dendritic spines. Nat Commun 7, 13282 (2016).

32. E.-M. Blumrich, J. C. Nicholson-Fish, M. Pronot, E. C. Davenport, D. Kurian, A. Cole, K. J. Smillie, M. A. Cousin, Phosphatidylinositol 4-kinase IIα is a glycogen synthase kinase 3-regulated interaction hub for activity-dependent bulk endocytosis. Cell Rep 42, 112633 (2023).

33. S. Wei, J. Zhang, R. Shi, Z. Yu, X. Chen, H. Wang, Identification of an integrated kinase-related prognostic gene signature associated with tumor immune microenvironment in human uterine corpus endometrial carcinoma. Front Oncol 12, 944000 (2022).

34. R. T. Sussman, J. L. Rokita, K. Huang, P. Raman, K. S. Rathi, D. Martinez, K. R. Bosse, M. Lane, L. S. Hart, T. Bhatti, B. Pawel, J. M. Maris, CAMKV Is a Candidate Immunotherapeutic Target in MYCN Amplified Neuroblastoma. Front Oncol 10, 302 (2020).

35. O. Shalem, N. E. Sanjana, E. Hartenian, X. Shi, D. A. Scott, T. Mikkelson, D. Heckl, B. L. Ebert, D. E. Root, J. G. Doench, F. Zhang, Genome-scale CRISPR-Cas9 knockout screening in human cells. Science 343, 84–87 (2014).

36. N. E. Sanjana, O. Shalem, F. Zhang, Improved vectors and genome-wide libraries for CRISPR screening. Nat Methods 11, 783–784 (2014).

37. Y. Fan, J. Cheng, S. A. Vasudevan, R. H. Patel, L. Liang, X. Xu, Y. Zhao, W. Jia, F. Lu, H. Zhang, J. G. Nuchtern, E. S. Kim, J. Yang, TAK1 inhibitor 5Z-7-oxozeaenol sensitizes neuroblastoma to chemotherapy. Apoptosis 18, 1224–1234 (2013).

38. S. Y. Jung, J. M. Choi, M. W. C. Rousseaux, A. Malovannaya, J. J. Kim, J. Kutzera, Y. Wang, Y. Huang, W. Zhu, S. Maity, H. Y. Zoghbi, J. Qin, An Anatomically Resolved Mouse Brain Proteome Reveals Parkinson Disease-relevant Pathways. Mol Cell Proteomics 16, 581–593 (2017).

39. M. Mirdita, K. Schütze, Y. Moriwaki, L. Heo, S. Ovchinnikov, M. Steinegger, ColabFold: making protein folding accessible to all. Nat Methods 19, 679–682 (2022).

40. H. M. Berman, J. Westbrook, Z. Feng, G. Gilliland, T. N. Bhat, H. Weissig, I. N. Shindyalov, P. E. Bourne, The Protein Data Bank. Nucleic Acids Res 28, 235–242 (2000).

41. J.-L. Wei, Z.-X. Fu, M. Fang, Q.-Y. Zhou, Q.-N. Zhao, J.-B. Guo, W.-D. Lu, H. Wang, High expression of CASK correlates with progression and poor prognosis of colorectal cancer. Tumour Biol 35, 9185–9194 (2014).

42. D. E. Frigo, M. K. Howe, B. M. Wittmann, A. M. Brunner, I. Cushman, Q. Wang, M. Brown, A. R. Means, D. P. McDonnell, CaM kinase kinase beta-mediated activation of the growth regulatory kinase AMPK is required for androgen-dependent migration of prostate cancer cells. Cancer Res 71, 528–537 (2011).

43. C. E. Massie, A. Lynch, A. Ramos-Montoya, J. Boren, R. Stark, L. Fazli, A. Warren, H. Scott, B. Madhu, N. Sharma, H. Bon, V. Zecchini, D.-M. Smith, G. M. Denicola, N. Mathews, M. Osborne, J. Hadfield, S. Macarthur, B. Adryan, S. K. Lyons, K. M. Brindle, J. Griffiths, M. E. Gleave, P. S. Rennie, D. E. Neal, I. G. Mills, The androgen receptor fuels prostate cancer by regulating central metabolism and biosynthesis. EMBO J 30, 2719–2733 (2011).

44. A. Bergamaschi, Y. H. Kim, K. A. Kwei, Y. La Choi, M. Bocanegra, A. Langerød, W. Han, D.-Y. Noh, D. G. Huntsman, S. S. Jeffrey, A.-L. Børresen-Dale, J. R. Pollack, CAMK1D amplification implicated in epithelial-mesenchymal transition in basal-like breast cancer. Mol Oncol 2, 327–339 (2008).

45. D. R. Alessi, Y. Saito, D. G. Campbell, P. Cohen, G. Sithanandam, U. Rapp, A. Ashworth, C. J. Marshall, S. Cowley, Identification of the sites in MAP kinase kinase-1 phosphorylated by p74raf-1. EMBO J 13, 1610–1619 (1994).

46. L. N. Johnson, M. E. Noble, D. J. Owen, Active and inactive protein kinases: structural basis for regulation. Cell 85, 149–158 (1996).

47. D. M. Payne, A. J. Rossomando, P. Martino, A. K. Erickson, J. H. Her, J. Shabanowitz, D. F. Hunt, M. J. Weber, T. W. Sturgill, Identification of the regulatory phosphorylation sites in pp42/mitogen-activated protein kinase (MAP kinase). EMBO J 10, 885–892 (1991).

48. M. Delhase, M. Hayakawa, Y. Chen, M. Karin, Positive and negative regulation of IkappaB kinase activity through IKKbeta subunit phosphorylation. Science 284, 309–313 (1999).

49. F. Mercurio, H. Zhu, B. W. Murray, A. Shevchenko, B. L. Bennett, J. Li, D. B. Young, M. Barbosa, M. Mann, A. Manning, A. Rao, IKK-1 and IKK-2: cytokine-activated IkappaB kinases essential for NF-kappaB activation. Science 278, 860–866 (1997).

50. D. M. Feliciano, A. M. Edelman, Repression of Ca2+/calmodulin-dependent protein kinase IV signaling accelerates retinoic acid-induced differentiation of human neuroblastoma cells. J Biol Chem 284, 26466–26481 (2009).

51. T. K. Blackwell, L. Kretzner, E. M. Blackwood, R. N. Eisenman, H. Weintraub, Sequence-specific DNA binding by the c-Myc protein. Science 250, 1149–1151 (1990).

52. E. M. Blackwood, R. N. Eisenman, Max: a helix-loop-helix zipper protein that forms a sequence-specific DNA-binding complex with Myc. Science 251, 1211–1217 (1991).

53. J. M. Shohet, R. Ghosh, C. Coarfa, A. Ludwig, A. L. Benham, Z. Chen, D. M. Patterson, E. Barbieri, P. Mestdagh, D. N. Sikorski, A. Milosavljevic, E. S. Kim, P. H. Gunaratne, A genome-wide search for promoters that respond to increased MYCN reveals both new oncogenic and tumor suppressor microRNAs associated with aggressive neuroblastoma. Cancer Res 71, 3841–3851 (2011).

54. N. J. Balamuth, A. Wood, Q. Wang, J. Jagannathan, P. Mayes, Z. Zhang, Z. Chen, E. Rappaport, J. Courtright, B. Pawel, B. Weber, R. Wooster, E. O. Sekyere, G. M. Marshall, J. M. Maris, Serial transcriptome analysis and cross-species integration identifies centromere-associated protein E as a novel neuroblastoma target. Cancer Res 70, 2749– 2758 (2010).

55. L. C. Heukamp, T. Thor, A. Schramm, K. De Preter, C. Kumps, B. De Wilde, A. Odersky, M. Peifer, S. Lindner, A. Spruessel, F. Pattyn, P. Mestdagh, B. Menten, S. Kuhfittig-Kulle, A. Künkele, K. König, L. Meder, S. Chatterjee, R. T. Ullrich, S. Schulte, J. Vandesompele, F. Speleman, R. Büttner, A. Eggert, J. H. Schulte, Targeted expression of mutated ALK induces neuroblastoma in transgenic mice. Sci Transl Med 4, 141ra91 (2012).

56. B. E. Lonze, D. D. Ginty, Function and regulation of CREB family transcription factors in the nervous system. Neuron 35, 605–623 (2002).

57. Z. Chen, C. Lei, C. Wang, N. Li, M. Srivastava, M. Tang, H. Zhang, J. M. Choi, S. Y. Jung, J. Qin, J. Chen, Global phosphoproteomic analysis reveals ARMC10 as an AMPK substrate that regulates mitochondrial dynamics. Nat Commun 10, 104 (2019).

58. J.-Y. Kim, E. A. Welsh, U. Oguz, B. Fang, Y. Bai, F. Kinose, C. Bronk, L. L. Remsing Rix, A. A. Beg, U. Rix, S. A. Eschrich, J. M. Koomen, E. B. Haura, Dissection of TBK1 signaling via phosphoproteomics in lung cancer cells. Proc Natl Acad Sci U S A 110, 12414–12419 (2013).

59. R. Dong, R. Yang, Y. Zhan, H.-D. Lai, C.-J. Ye, X.-Y. Yao, W.-Q. Luo, X.-M. Cheng, J.-J. Miao, J.-F. Wang, B.-H. Liu, X.-Q. Liu, L.-L. Xie, Y. Li, M. Zhang, L. Chen, W.-C. Song, W. Qian, W.-Q. Gao, Y.-H. Tang, C.-Y. Shen, W. Jiang, G. Chen, W. Yao, K.-R. Dong, X.-M. Xiao, S. Zheng, K. Li, J. Wang, Single-Cell Characterization of Malignant Phenotypes and Developmental Trajectories of Adrenal Neuroblastoma. Cancer Cell 38, 716–733.e6 (2020).

60. O. M. Enache, V. Rendo, M. Abdusamad, D. Lam, D. Davison, S. Pal, N. Currimjee, J. Hess, S. Pantel, A. Nag, A. R. Thorner, J. G. Doench, F. Vazquez, R. Beroukhim, T. R. Golub, U. Ben-David, Cas9 activates the p53 pathway and selects for p53-inactivating mutations. Nat Genet 52, 662–668 (2020).

61. Y. Fu, J. A. Foden, C. Khayter, M. L. Maeder, D. Reyon, J. K. Joung, J. D. Sander, High-frequency off-target mutagenesis induced by CRISPR-Cas nucleases in human cells. Nat Biotechnol 31, 822–826 (2013).

62. P. D. Hsu, D. A. Scott, J. A. Weinstein, F. A. Ran, S. Konermann, V. Agarwala, Y. Li, E. J. Fine, X. Wu, O. Shalem, T. J. Cradick, L. A. Marraffini, G. Bao, F. Zhang, DNA targeting specificity of RNA-guided Cas9 nucleases. Nat Biotechnol 31, 827–832 (2013).

63. A. Li, M. R. Tanner, C. M. Lee, A. E. Hurley, M. De Giorgi, K. E. Jarrett, T. H. Davis, A. M. Doerfler, G. Bao, C. Beeton, W. R. Lagor, AAV-CRISPR Gene Editing Is Negated by Pre-existing Immunity to Cas9. Mol Ther 28, 1432–1441 (2020).

64. J. Cao, L. Wu, S.-M. Zhang, M. Lu, W. K. C. Cheung, W. Cai, M. Gale, Q. Xu, Q. Yan, An easy and efficient inducible CRISPR/Cas9 platform with improved specificity for multiple gene targeting. Nucleic Acids Res 44, e149 (2016).

65. F. González, Z. Zhu, Z.-D. Shi, K. Lelli, N. Verma, Q. V. Li, D. Huangfu, An iCRISPR platform for rapid, multiplexable, and inducible genome editing in human pluripotent stem cells. Cell Stem Cell 15, 215–226 (2014).

66. L. E. Dow, J. Fisher, K. P. O’Rourke, A. Muley, E. R. Kastenhuber, G. Livshits, D. F. Tschaharganeh, N. D. Socci, S. W. Lowe, Inducible in vivo genome editing with CRISPR-Cas9. Nat Biotechnol 33, 390–394 (2015).

67. B. J. Aubrey, G. L. Kelly, A. J. Kueh, M. S. Brennan, L. O’Connor, L. Milla, S. Wilcox, L. Tai, A. Strasser, M. J. Herold, An inducible lentiviral guide RNA platform enables the identification of tumor-essential genes and tumor-promoting mutations in vivo. Cell Rep 10, 1422–1432 (2015).

68. S. Guan, J. Lu, Y. Zhao, Y. Yu, H. Li, Z. Chen, Z. Shi, H. Liang, M. Wang, K. Guo, X. Chen, W. Sun, S. Bieerkehazhi, X. Xu, S. Sun, S. Agarwal, J. Yang, MELK is a novel therapeutic target in high-risk neuroblastoma. Oncotarget 9, 2591–2602 (2017).

69. A. Chlenski, C. Park, M. Dobratic, H. R. Salwen, B. Budke, J.-H. Park, R. Miller, M. A. Applebaum, E. Wilkinson, Y. Nakamura, P. P. Connell, S. L. Cohn, Maternal Embryonic Leucine Zipper Kinase (MELK), a Potential Therapeutic Target for Neuroblastoma. Mol Cancer Ther 18, 507–516 (2019).

70. P. D. Marley, K. A. Thomson, The Ca++/calmodulin-dependent protein kinase II inhibitors KN62 and KN93, and their inactive analogues KN04 and KN92, inhibit nicotinic activation of tyrosine hydroxylase in bovine chromaffin cells. Biochem Biophys Res Commun 221, 15–18 (1996).

71. H. Tokumitsu, H. Inuzuka, Y. Ishikawa, M. Ikeda, I. Saji, R. Kobayashi, STO-609, a specific inhibitor of the Ca(2+)/calmodulin-dependent protein kinase kinase. J Biol Chem 277, 15813–15818 (2002).

72. B. Knüsel, F. Hefti, K-252 compounds: modulators of neurotrophin signal transduction. J Neurochem 59, 1987–1996 (1992).

73. S. Fabre, M. Prudhomme, M. Rapp, Protein kinase C inhibitors; structure-activity relationships in K252c-related compounds. Bioorg Med Chem 1, 193–196 (1993).

74. J. Meiler, M. Guyot, S. Hoffarth, E. Wesarg, Y. Höhn, F. Breitenbuecher, M. Schuler, Individual dose and scheduling determine the efficacy of combining cytotoxic anticancer agents with a kinase inhibitor in non-small-cell lung cancer. J Cancer Res Clin Oncol 138, 1385–1394 (2012).

75. T. Tamaoki, H. Nomoto, I. Takahashi, Y. Kato, M. Morimoto, F. Tomita, Staurosporine, a potent inhibitor of phospholipid/Ca++dependent protein kinase. Biochem Biophys Res Commun 135, 397–402 (1986).

76. B. B. Touré, J. Giraldes, T. Smith, E. R. Sprague, Y. Wang, S. Mathieu, Z. Chen, Y. Mishina, Y. Feng, Y. Yan-Neale, S. Shakya, D. Chen, M. Meyer, D. Puleo, J. T. Brazell, C. Straub, D. Sage, K. Wright, Y. Yuan, X. Chen, J. Duca, S. Kim, L. Tian, E. Martin, K. Hurov, W. Shao, Toward the Validation of Maternal Embryonic Leucine Zipper Kinase: Discovery, Optimization of Highly Potent and Selective Inhibitors, and Preliminary Biology Insight. J Med Chem 59, 4711–4723 (2016).

77. L. Li, A. C. Hung, A. G. Porter, Secretogranin II: a key AP-1-regulated protein that mediates neuronal differentiation and protection from nitric oxide-induced apoptosis of neuroblastoma cells. Cell Death Differ 15, 879–888 (2008).

78. S. Scala, K. Wosikowski, P. Giannakakou, P. Valle, J. L. Biedler, B. A. Spengler, E. Lucarelli, S. E. Bates, C. J. Thiele, Brain-derived neurotrophic factor protects neuroblastoma cells from vinblastine toxicity. Cancer Res 56, 3737–3742 (1996).

79. D. S. Middlemas, B. K. Kihl, J. Zhou, X. Zhu, Brain-derived neurotrophic factor promotes survival and chemoprotection of human neuroblastoma cells. J Biol Chem 274, 16451– 16460 (1999).

80. D. P. Radin, P. Patel, BDNF: An Oncogene or Tumor Suppressor? Anticancer Res 37, 3983–3990 (2017).

81. C. M. Schworer, R. J. Colbran, T. R. Soderling, Reversible generation of a Ca2+- independent form of Ca2+(calmodulin)-dependent protein kinase II by an autophosphorylation mechanism. J Biol Chem 261, 8581–8584 (1986).

82. Y. Hashimoto, C. M. Schworer, R. J. Colbran, T. R. Soderling, Autophosphorylation of Ca2+/calmodulin-dependent protein kinase II. Effects on total and Ca2+-independent activities and kinetic parameters. J Biol Chem 262, 8051–8055 (1987).

83. J. M. Johnson, J. Castle, P. Garrett-Engele, Z. Kan, P. M. Loerch, C. D. Armour, R. Santos, E. E. Schadt, R. Stoughton, D. D. Shoemaker, Genome-wide survey of human alternative pre-mRNA splicing with exon junction microarrays. Science 302, 2141–2144 (2003).

84. E. S. Lander, L. M. Linton, B. Birren, C. Nusbaum, M. C. Zody, J. Baldwin, K. Devon, K. Dewar, M. Doyle, W. FitzHugh, R. Funke, D. Gage, K. Harris, A. Heaford, J. Howland, L. Kann, J. Lehoczky, R. LeVine, P. McEwan, K. McKernan, J. Meldrim, J. P. Mesirov, C. Miranda, W. Morris, J. Naylor, C. Raymond, M. Rosetti, R. Santos, A. Sheridan, C. Sougnez, Y. Stange-Thomann, N. Stojanovic, A. Subramanian, D. Wyman, J. Rogers, J. Sulston, R. Ainscough, S. Beck, D. Bentley, J. Burton, C. Clee, N. Carter, A. Coulson, R. Deadman, P. Deloukas, A. Dunham, I. Dunham, R. Durbin, L. French, D. Grafham, S. Gregory, T. Hubbard, S. Humphray, A. Hunt, M. Jones, C. Lloyd, A. McMurray, L. Matthews, S. Mercer, S. Milne, J. C. Mullikin, A. Mungall, R. Plumb, M. Ross, R. Shownkeen, S. Sims, R. H. Waterston, R. K. Wilson, L. W. Hillier, J. D. McPherson, M. A. Marra, E. R. Mardis, L. A. Fulton, A. T. Chinwalla, K. H. Pepin, W. R. Gish, S. L. Chissoe, M. C. Wendl, K. D. Delehaunty, T. L. Miner, A. Delehaunty, J. B. Kramer, L. L. Cook, R. S. Fulton, D. L. Johnson, P. J. Minx, S. W. Clifton, T. Hawkins, E. Branscomb, P. Predki, P. Richardson, S. Wenning, T. Slezak, N. Doggett, J. F. Cheng, A. Olsen, S. Lucas, C. Elkin, E. Uberbacher, M. Frazier, R. A. Gibbs, D. M. Muzny, S. E. Scherer, J. B. Bouck, E. J. Sodergren, K. C. Worley, C. M. Rives, J. H. Gorrell, M. L. Metzker, S. L. Naylor, R. S. Kucherlapati, D. L. Nelson, G. M. Weinstock, Y. Sakaki, A. Fujiyama, M. Hattori, T. Yada, A. Toyoda, T. Itoh, C. Kawagoe, H. Watanabe, Y. Totoki, T. Taylor, J. Weissenbach, R. Heilig, W. Saurin, F. Artiguenave, P. Brottier, T. Bruls, E. Pelletier, C. Robert, P. Wincker, D. R. Smith, L. Doucette-Stamm, M. Rubenfield, K. Weinstock, H. M. Lee, J. Dubois, A. Rosenthal, M. Platzer, G. Nyakatura, S. Taudien, A. Rump, H. Yang, J. Yu, J. Wang, G. Huang, J. Gu, L. Hood, L. Rowen, A. Madan, S. Qin, R. W. Davis, N. A. Federspiel, A. P. Abola, M. J. Proctor, R. M. Myers, J. Schmutz, M. Dickson, J. Grimwood, D. R. Cox, M. V. Olson, R. Kaul, C. Raymond, N. Shimizu, K. Kawasaki, S. Minoshima, G. A. Evans, M. Athanasiou, R. Schultz, B. A. Roe, F. Chen, H. Pan, J. Ramser, H. Lehrach, R. Reinhardt, W. R. McCombie, M. de la Bastide, N. Dedhia, H. Blöcker, K. Hornischer, G. Nordsiek, R. Agarwala, L. Aravind, J. A. Bailey, A. Bateman, S. Batzoglou, E. Birney, P. Bork, D. G. Brown, C. B. Burge, L. Cerutti, H. C. Chen, D. Church, M. Clamp, R. R. Copley, T. Doerks, S. R. Eddy, E. E. Eichler, T. S. Furey, J. Galagan, J. G. Gilbert, C. Harmon, Y. Hayashizaki, D. Haussler, H. Hermjakob, K. Hokamp, W. Jang, L. S. Johnson, T. A. Jones, S. Kasif, A. Kaspryzk, S. Kennedy, W. J. Kent, P. Kitts, E. V. Koonin, I. Korf, D. Kulp, D. Lancet, T. M. Lowe, A. McLysaght, T. Mikkelsen, J. V. Moran, N. Mulder, V. J. Pollara, C. P. Ponting, G. Schuler, J. Schultz, G. Slater, A. F. Smit, E. Stupka, J. Szustakowki, D. Thierry-Mieg, J. Thierry-Mieg, L. Wagner, J. Wallis, R. Wheeler, A. Williams, Y. I. Wolf, K. H. Wolfe, S. P. Yang, R. F. Yeh, F. Collins, M. S. Guyer, J. Peterson, A. Felsenfeld, K. A. Wetterstrand, A. Patrinos, M. J. Morgan, P. de Jong, J. J. Catanese, K. Osoegawa, H. Shizuya, S. Choi, Y. J. Chen, J. Szustakowki, International Human Genome Sequencing Consortium, Initial sequencing and analysis of the human genome. Nature 409, 860–921 (2001).

85. T. W. Nilsen, B. R. Graveley, Expansion of the eukaryotic proteome by alternative splicing. Nature 463, 457–463 (2010).

86. X. Guo, Q.-R. Chen, Y. K. Song, J. S. Wei, J. Khan, Exon array analysis reveals neuroblastoma tumors have distinct alternative splicing patterns according to stage and MYCN amplification status. BMC Med Genomics 4, 35 (2011).

87. J. Chen, C. S. Hackett, S. Zhang, Y. K. Song, R. J. A. Bell, A. M. Molinaro, D. A. Quigley, A. Balmain, J. S. Song, J. F. Costello, W. C. Gustafson, T. Van Dyke, P.-Y. Kwok, J. Khan, W. A. Weiss, The genetics of splicing in neuroblastoma. Cancer Discov 5, 380–395 (2015).

88. S. Zhang, J. S. Wei, S. Q. Li, T. C. Badgett, Y. K. Song, S. Agarwal, C. Coarfa, C. Tolman, L. Hurd, H. Liao, J. He, X. Wen, Z. Liu, C. J. Thiele, F. Westermann, S. Asgharzadeh, R. C. Seeger, J. M. Maris, J. M. Guidry Auvil, M. A. Smith, E. D. Kolaczyk, J. Shohet, J. Khan, MYCN controls an alternative RNA splicing program in high-risk metastatic neuroblastoma. Cancer Lett 371, 214–224 (2016).

89. K. R. Bosse, S. J. Diskin, K. A. Cole, A. C. Wood, R. W. Schnepp, G. Norris, L. B. Nguyen, J. Jagannathan, M. Laquaglia, C. Winter, M. Diamond, C. Hou, E. F. Attiyeh, Y. P. Mosse, V. Pineros, E. Dizin, Y. Zhang, S. Asgharzadeh, R. C. Seeger, M. Capasso, B. R. Pawel, M. Devoto, H. Hakonarson, E. F. Rappaport, I. Irminger-Finger, J. M. Maris, Common variation at BARD1 results in the expression of an oncogenic isoform that influences neuroblastoma susceptibility and oncogenicity. Cancer Res 72, 2068–2078 (2012).

90. A. Hudmon, H. Schulman, Neuronal CA2+/calmodulin-dependent protein kinase II: the role of structure and autoregulation in cellular function. Annu Rev Biochem 71, 473–510 (2002).

91. K. U. Bayer, P. De Koninck, H. Schulman, Alternative splicing modulates the frequency-dependent response of CaMKII to Ca(2+) oscillations. EMBO J 21, 3590–3597 (2002).

92. S. A. Walter, R. E. Cutler, R. Martinez, M. Gishizky, R. J. Hill, Stk10, a new member of the polo-like kinase kinase family highly expressed in hematopoietic tissue. J Biol Chem 278, 18221–18228 (2003).

93. R. Viswanatha, P. Y. Ohouo, M. B. Smolka, A. Bretscher, Local phosphocycling mediated by LOK/SLK restricts ezrin function to the apical aspect of epithelial cells. J Cell Biol 199, 969–984 (2012).

94. S. Arora, I. M. Gonzales, R. T. Hagelstrom, C. Beaudry, A. Choudhary, C. Sima, R. Tibes, S. Mousses, D. O. Azorsa, RNAi phenotype profiling of kinases identifies potential therapeutic targets in Ewing’s sarcoma. Mol Cancer 9, 218 (2010).

95. F. Weinberg, N. Reischmann, L. Fauth, S. Taromi, J. Mastroianni, M. Köhler, S. Halbach, A. C. Becker, N. Deng, T. Schmitz, F. M. Uhl, N. Herbener, B. Riedel, F. Beier, A. Swarbrick, S. Lassmann, J. Dengjel, R. Zeiser, T. Brummer, The Atypical Kinase RIOK1 Promotes Tumor Growth and Invasive Behavior. EBioMedicine 20, 79–97 (2017).

96. H. Kase, K. Iwahashi, Y. Matsuda, K-252a, a potent inhibitor of protein kinase C from microbial origin. J Antibiot (Tokyo) 39, 1059–1065 (1986).

97. H. Kase, K. Iwahashi, S. Nakanishi, Y. Matsuda, K. Yamada, M. Takahashi, C. Murakata, A. Sato, M. Kaneko, K-252 compounds, novel and potent inhibitors of protein kinase C and cyclic nucleotide-dependent protein kinases. Biochem Biophys Res Commun 142, 436–440 (1987).

98. Y. Hashimoto, T. Nakayama, T. Teramoto, H. Kato, T. Watanabe, M. Kinoshita, K. Tsukamoto, K. Tokunaga, K. Kurokawa, S. Nakanishi, Potent and preferential inhibition of Ca2+/calmodulin-dependent protein kinase II by K252a and its derivative, KT5926. Biochem Biophys Res Commun 181, 423–429 (1991).

99. S. Koizumi, M. L. Contreras, Y. Matsuda, T. Hama, P. Lazarovici, G. Guroff, K-252a: a specific inhibitor of the action of nerve growth factor on PC 12 cells. J Neurosci 8, 715–721 (1988).

100. P. Tapley, F. Lamballe, M. Barbacid, K252a is a selective inhibitor of the tyrosine protein kinase activity of the trk family of oncogenes and neurotrophin receptors. Oncogene 7, 371–381 (1992).

101. S. H. Nye, S. P. Squinto, D. J. Glass, T. N. Stitt, P. Hantzopoulos, M. J. Macchi, N. S. Lindsay, N. Y. Ip, G. D. Yancopoulos, K-252a and staurosporine selectively block autophosphorylation of neurotrophin receptors and neurotrophin-mediated responses. Mol Biol Cell 3, 677–686 (1992).

