## Supplemental Table 2 for "CAMKV Kinase Signaling Is a Novel Therapeutic Avenue with Prognostic Relevance in Neuroblastoma"

Table S2. Target sequences of gene knockdown

| Target gene | sequence |
| --- | --- |
| sh-CREB-1 | aagcagctcgagagtgtcgta |
| sh-CREB-2 | aagctgcctctggagacgtac |
| sh-CCNA2-1 | aactggatcaatttgctgact |
| sh-CCNA2-2 | aacccaccagagacactaaat |
| sh-CAMKV-1 | Aacctggcaaataaacatcac |
| sh-CAMKV-2 | Aacctcatgagcctctaaggg |
| sh-MYCN-1 | aattcttacactgcctgtata |
| sh-MYCN-2 | aatctctgttatgtactgtac |
| sh-MYC-1 | aatgtcctgagcaatcaccta |
| sh-MYC-2 | aatgcatgatcaaatgcaacc |
| sh-Control | CTGGCATCGGTGTGGATGA |
