## Supplemental Table 3 for "CAMKV Kinase Signaling Is a Novel Therapeutic Avenue with Prognostic Relevance in Neuroblastoma"

Table S3. Target sequences of CRISPR-Cas9 gene knockout

| Gene | Target sequence 1 | Target sequence 2 |
| --- | --- | --- |
| CAMK1 | GGAGCTCTTTGACCGTATTG | GATCCCGGTGTACAATGCCC |
| CAMK1D | AGAGCTGTTTGACCGGATAG | AGTACACGGCGTCCAAGACT |
| CAMK1G | CTGTGTAGACACCCCGCTCC | GGACGATGCCATTCTCATGT |
| CAMK2A | GGAACTGTTTGAAGATATCG | CCGGGAGTATTACAGTGAGG |
| CAMK2B | CTCTGTGGACGACCCCCATT | GACAATGGAGAACGGCCTCC |
| CAMK2D | GCTTAGCCATAGAAGTTCAA | GCTTTTAGCTAGCAAATCCA |
| CAMK2G | CCACTGTATACATCAGATTC | ACCAGCATGACATCGTCCAC |
| CAMK4 | TCTCTGGTTTGAGATCACGA | CGATTTTGAGTGGTGCATCT |
| CAMKV | CAACGTGGTACGGCAAGTCC | ACTCACTCAAGATCGTGCAC |
| CASK | CCATTATATGAGACAGATAC | ATGATAATAACATAATTCAC |
| PNCK | CGAGCTGTTTGACCGCATCA | GCGCGGCTCCTACACAGAGA |
