## Supplemental Table 4 for "CAMKV Kinase Signaling Is a Novel Therapeutic Avenue with Prognostic Relevance in Neuroblastoma"

Table S4. Primer sequences

| Gene | Forward primer | Reverse primer |
| --- | --- | --- |
| CAMKV | TGCAGAAGAGGCCATCTCCC | AGCCGTTTCATGAGGGTGGT |
| CCNA1 | ACAGGAGGACCTGTGGCCAG | TCTTGGACCCCACAGTCAG |
| CCNA2 | GGACCCAGAAAACCATTGGT | TCATGGTAGTCTGGTACTTC |
| CCNB1 | GCCAGAACCTGAGCCAGAAC | TCAGAGAAAGCCTGACACAG |
| CCND1 | CAATGACCCCGCACGATTTC | CATGGAGGGCGGATTGGAA |
| CCNE1 | TTCTGGATTGGTTAATGGAG | ATGAAATCCCAATAAGCTG |
| MYCN | AGAGGACACCCTGAGCGATTC | CATAGTTGTGCTGCTGGTGGA |
| MYC | CTCCATGAGGAGACACCGCCCA | AAGGTGATCCAGACTCTGACCT |
| ACTB | TCTACAATGAGCTGCGTG | GATAGCACAGCCTGGATAG |
