## Supplemental Table 6 for "CAMKV Kinase Signaling Is a Novel Therapeutic Avenue with Prognostic Relevance in Neuroblastoma"

Table S6. CREB target genes that decreased in CAMKV knockdown cells

| Gene symbol | Official Full Name | Log2(Fold change) |
| --- | --- | --- |
| SCG2 | secretogranin II | -1.71 |
| BDNF | brain derived neurotrophic factor | -1.57 |
| NEFL | neurofilament light chain | -1.13 |
| TGFB2 | transforming growth factor beta 2 | -1.08 |
| ENO2 | enolase 2 | -0.92 |
| NF1 | neurofibromin 1 | -0.80 |
| Per2 | period circadian regulator 2 | -0.73 |
| ADCYAP1 | adenylate cyclase activating polypeptide 1 | -0.62 |
| SSTR2 | somatostatin receptor 2 | -0.62 |
| ATP1A1 | ATPase Na+/K+ transporting subunit alpha 1 | -0.59 |
| IGF1 | insulin like growth factor 1 | -0.57 |
